# Physical basis for the interaction between *Drosophila* ROS1 and the GPCR BOSS

**DOI:** 10.1101/2024.03.13.584817

**Authors:** Jianan Zhang, Yuko Tsutsui, Hengyi Li, Tongqing Li, Yueyue Wang, Daryl E. Klein

## Abstract

*Drosophila* ROS1 (dROS1, Sevenless) is a receptor tyrosine kinase (RTK) essential for the differentiation of *Drosophila* R7 photoreceptor cells*^1, 2^*. Activation of dROS1 is mediated by binding to the extracellular region (ECR) of the GPCR (G protein coupled receptor) BOSS (Bride Of Sevenless) on adjacent cells*^1, 3, 4^*. Genetic evidence together with *in vitro* activity assays confirmed the activation of dROS1 by BOSS and identified subsequent downstream signaling pathways including SOS (Son of Sevenless)*^1, 5^*. However, the physical basis for how dROS1 interacts with the GPCR BOSS has long remained unknown. Here we provide the first structure, using Cryo-Electron Microscopy (CryoEM), of dROS1’s extracellular region, which mediates ligand binding. We show that the N-terminal region of dROS1 adopts a folded-over conformation harboring a novel structural domain. We further narrowed down the interacting binding epitopes on both dROS1 and BOSS. This includes a beta-strand in dROS1’s third Fibronectin type III (FNIII) domain and the C-terminal portion of BOSS’ ECR. Our mutagenesis studies, coupled with AlphaFold complex predictions, support a binding interaction mediated by a hydrophobic interaction and beta-strand augmentation between these regions. Our findings provide a fundamental understanding of the regulatory function of dROS1 and further provide mechanistic insight into the human ortholog and oncogene ROS1.

## Introduction

*Drosophila* ROS1 (dROS1), also known as Sevenless, is a critical RTK required for R7 precursor cells to receive positional signals for the specification of R7 photoreceptors*^6^*. Specification of all other photoreceptor cells (R1-R6) require another RTK, *Drosophila* EGF receptor (dEGFR) – while dROS1 is selectively required for the differentiation of R7 cells*^7, 8^*. Interestingly, rather than a soluble morphogen or growth factor, the ligand for dROS1 is a membrane-bound GPCR, which is an unusual ligand for an RTK*^3^*. The first reported physical interaction between dROS1 and BOSS was a heterotypic cell-aggregation assay involving dROS1-expressing and BOSS-expressing S2 cells, and further supported by co-immunoprecipitation*^3, 9^*. It has been proposed that the positional proximity of dROS1 on noncommitted R7 precursor cells with BOSS on differentiated R8 cells promotes binding through cell-cell contacts*^10^*. However, no quantitative *in vitro* study has been performed to confirm nor has any structural study been done to shed light on the details of this unusual interaction.

Like all other RTKs, the architecture of dROS1 is composed of an extracellular region (ECR), a single-pass transmembrane domain (TM), and an intracellular kinase domain (KD). Notably, ROS1 has the largest ECR of any described RTK. Human ROS1 and dROS1 both are predicted to have an ECR composed of nine fibronectin type III domains (FNIII) and three YWTD beta-propellers **(Figure 1a)**. Importantly, however, no structural studies have been done on ROS1 from any species to date. Therefore, there is no experimental evidence for the ECR architecture of ROS1. Interestingly, dROS1 contains a poly-arginine (9 consecutive arginines) sequence in the last FNIII domain just prior to the transmembrane domain. This sequence harbors multiple consensus furin proteolytic cleavage sites (i.e. RXRR) **(Figure 1a)**. Notably, cleavage occurs topologically in the same FNIII loop where insulin receptor is processed*^11^*. Previous literature suggests that the large ECR of dROS1 is cleaved at this position but remains non-covalently associated with the rest of the receptor*^12^*. This is in contrast to the insulin receptor which is cleaved but remains covalently associated through disulfide bonds to the kinase domain*^13, 14^*.

**Fig. 1.**
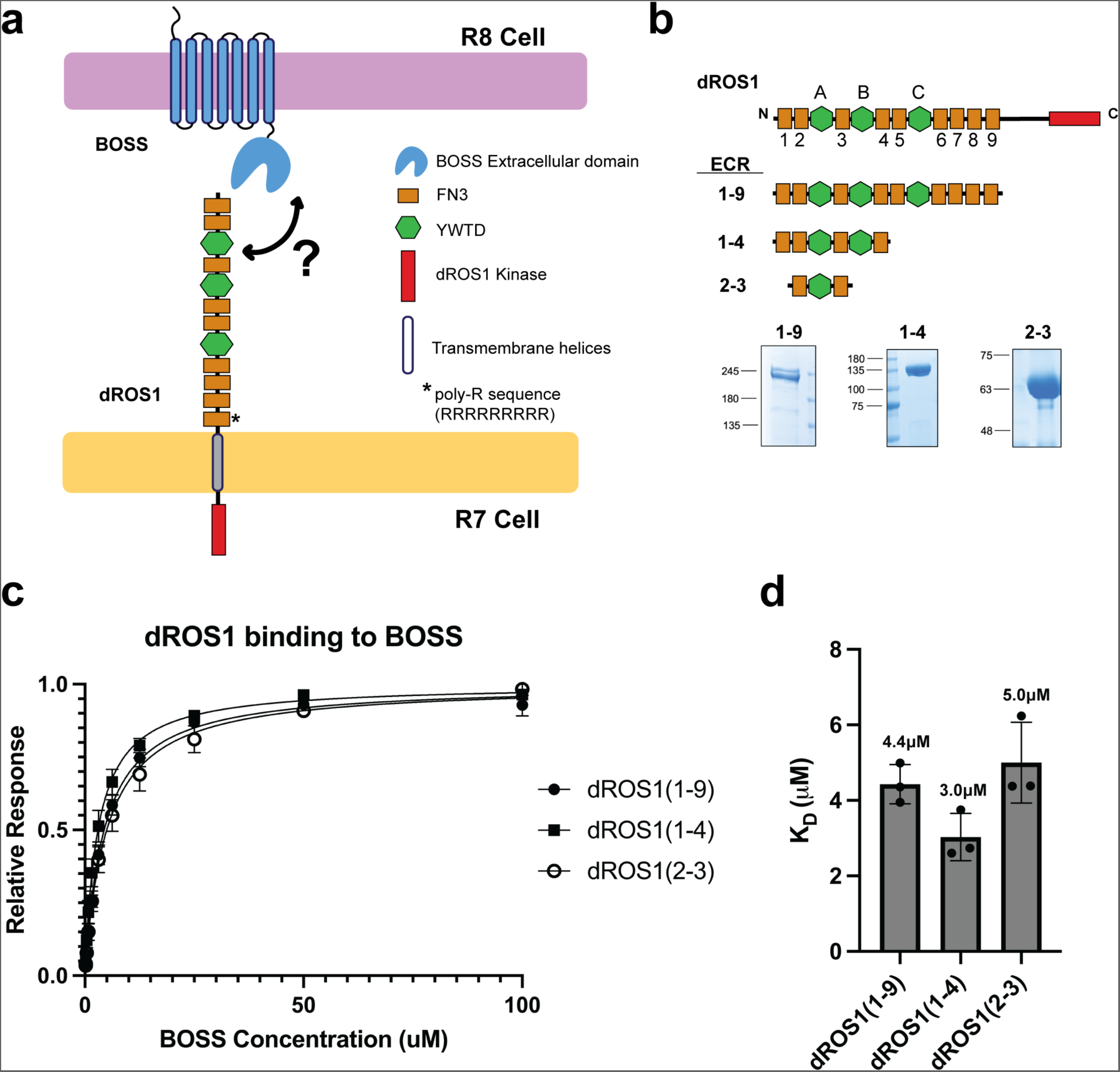
The N-terminal region of dROS1 interacts with BOSS’ ECR. **a**, Illustration of dROS1 and BOSS architecture. The large extracellular region of dROS1 is composed of 9 Fibronectin type III (FNIII) domains (number 1-9, orange rectangles) and 3 YWTD propellers (A-C, green hexagons). BOSS is a GPCR with an extracellular region and 7 transmembrane helices. The binding region of dROS1 to BOSS remains unknown. **b**, Domain organization and ECR, name designation based upon terminal FNIII domains, and representative Coomassie stained SDS-page gel analysis of different dROS1 ECR constructs. **c**, Biacore T200 binding analysis of dROS1’s ECR constructs to BOSS’ ECR. The minimal construct of dROS1 (2-3, second to third FNIII domain) binds to BOSS as well as the full ECR (n=3 for all three constructs). **d**, Summary of K_D_ of all tested constructs. One-way ANOVA test shows no significant difference among means (P > 0.05).

Surprisingly, the soluble ECR of BOSS is insufficient to activate – and paradoxically inhibits – dROS1 even when presented as a preformed oligomer*^9^*. This is contrary to the ‘central dogma’ of RTK oligomer-induced activation*^15, 16^*. However, a dROS1 mutant that removes most of the ECR is constitutively active*^17^*. In addition, overexpression of dROS1 or a chimeric dROS1 with the kinase domain of dEGFR (Drosophila Epidermal Growth Factor Receptor) shows no activation in the absence of ligand, revealing a potent inhibitory role of dROS1’s ECR*^6^*. Together, this points to a unique regulatory mechanism employed by dROS1 unlike that of other well-studied RTKs. To further investigate this activation mechanism, we solved the structure of dROS1’s ECR and analyzed the biophysical basis for the dROS1-BOSS interaction.

## Results

### The N-terminal region of dROS1 interacts with the GPCR BOSS

To characterize the binding interaction between dROS1 and BOSS, we purified the full-length ECR of dROS1 (construct 1-9 in **Figure 1b**) – as well as multiple smaller ECR fragments **(Figure 1b)** – and examined their binding affinities towards the ECR of BOSS using surface plasmon resonance (SPR, Biacore) **(Figure 1c, 1d)**. We found that the full ECR of dROS1 binds to BOSS with a K_D_ of 4.4 µM **(Figure 1c, 1d)**. Unlike other RTKs with soluble ligands (i.e. hormones and morphogens) that typically demonstrate nanomolar affinity, dROS1 binds to BOSS at a comparatively modest binding (µM) affinity. As BOSS is also a membrane-bound receptor, this binding affinity is consistent with other cell-cell adhesion receptors (i.e. notch and delta*^18^*). We then determined that the N-terminal region of dROS1 (constructs 1-4 and 2-3) also binds to BOSS with a similar affinity **(Figure 1c, 1d)**. Thus, these truncations narrow down the binding epitope to the region flanked by the second and third FNIII domains (construct 2-3), and include the first YWTD propeller domain.

### The N-terminal region of dROS1 adopts a folded-over conformation

To establish a biophysical understanding of dROS1 function we sought to obtain a 3-Dimensional structure of dROS1’s ECR. Notably, there is no published structural information on the ECR of ROS1 from any species. We expressed and purified the dROS1 ECR (construct 1-6) and obtained a 3.94Å structure using single-particle CryoEM **(Figure 2a; Supplemental Figure 1)**. Surprisingly, model building only resolves the N-terminal region (equivalent to construct 1-4) starting from the N-terminus to the fourth FNIII domain. The C-terminal domains starting from the fifth FNIII domain to the sixth, including the intervening YWTD-C, were not observed in the map **(Figure 2a, 2b)**. We believe these C-terminal domains are flexible with respect to the N-terminal domains and thus not resolved in our reconstruction. The poor density and low resolution of the fourth FNIII domain also supports this idea **(Supplement Figure 1)**. Despite the flexibility, we were able to further improve the density of the fourth FNIII domain by masked local refinement **(Supplement Figure 2)**.

**Fig. 2.**
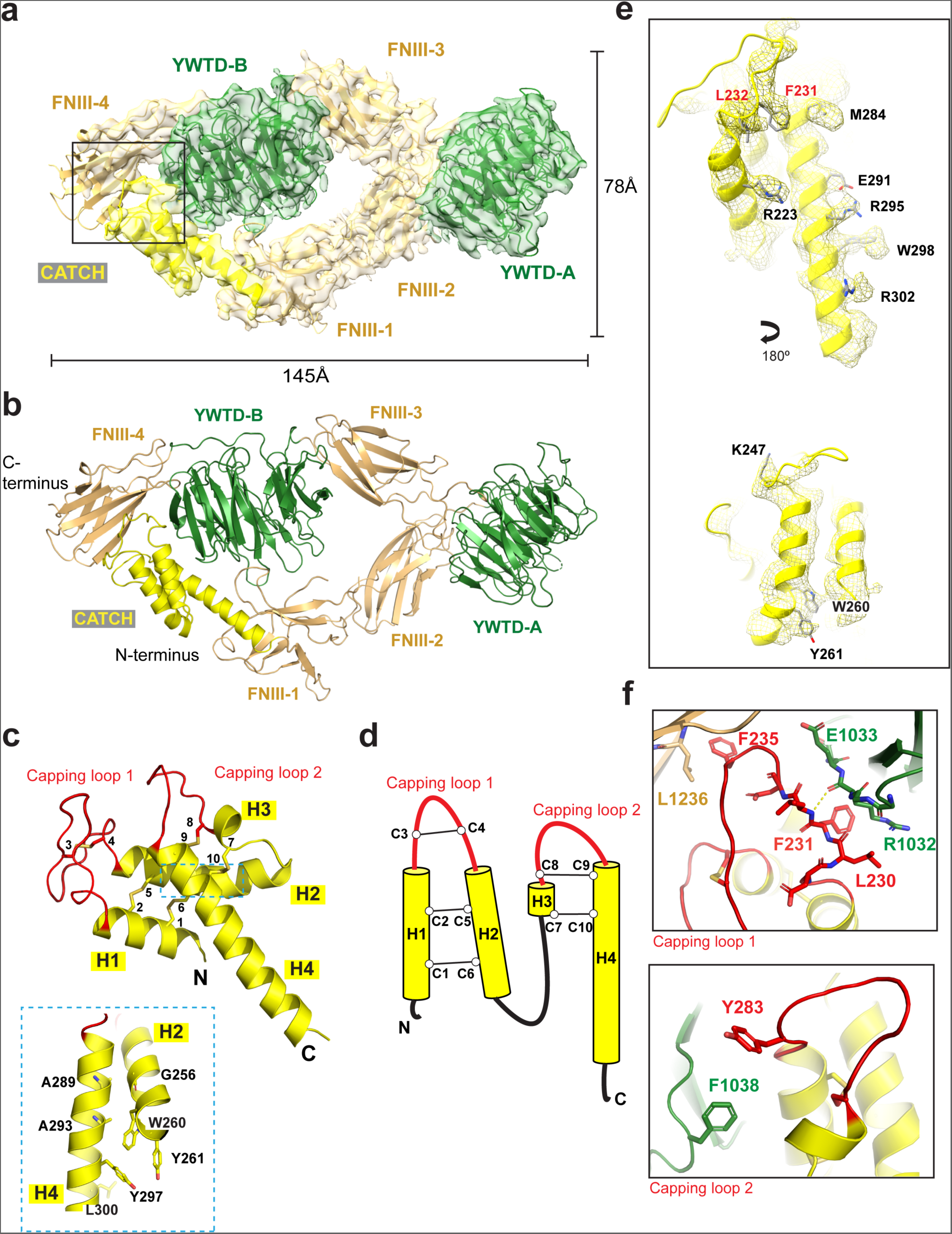
CryoEM Structure of the N-terminal ligand binding region of dROS1. **a**, Cartoon representation of dROS1 ECR structure and fitted into CryoEM map at 3.94Å resolution, with FNIII domains in light brown and YWTD domains in green. Only the N-terminal domains (construct 1-4) of dROS1’s ECR can be resolved in the reconstructed map. The N-terminal ECR of dROS1 adopts a folded-over conformation held by the interaction of the CATCH domain (colored in yellow) with the second propeller (YWTD-B) and fourth FNIII domain (FNIII-4). **b**, Cartoon representation of dROS1’s ECR structure. **c**, Structure of the CATCH domain. Capping loops colored in red. Cysteine residues are numbered from the N-terminus (1-10). Helices are labeled as H1-H4 (helix-1 to helix-4). H2 and H4 are associated through hydrophobic interactions through aromatic residues (W260, Y261, Y297 and L300). **d**, Illustration of the disulfide linkage in the CATCH domain. Disulfide bonds are formed between C1-C6, C2-C5 and C3-C4, of H1 and H2; C8-C9 and C7-C10 bridge H3 and H4. **e**, EM density maps of the CATCH domain. Despite the modest resolution, features of bulky side chains can be clearly observed. F231 and L232 colored in red are the capping residues as shown in **f**. **f**, Zoomed-in view of the capping interaction. The hydrophobic residue F235 in capping loop 1 is interacting with L1236 in FNIII-4. Same for the interaction with YWTD-B. The aromatic residue Y283 in capping loop 2 is interacting with F1038 from YWTD-B.

Interestingly, the model reveals that instead of being linearized, the N-terminal region adopts a folded-over conformation stabilized by an interaction of an N-terminal cysteine-rich region with the second propeller (YWTD-B) and fourth FNIII domain **(Figure 2a, 2b)**. The N-terminal cysteine-rich peptide forms a previously unappreciated domain. The domain is comprised of two disulfide stapled helical hairpins that associate with one another through hydrophobic interactions **(Figure 2c, 2d)**. The two helical hairpins generate two loops (capping loops 1 and 2) which stabilize the N-terminal ECR compacted fold. Despite the rather limited resolution in this region, we observe clear density for bulky side-chains which assisted in model building **(Figure 2e; Supplement Figure 3)**. In capping loop 1, F235 interacts with L1236 of the fourth FNIII domain. Additionally, L230 and F231 interact with the second YWTD propeller by hydrophobic contacts. This is further supported by potential hydrogen bonding to the main chain. Capping loop 2 interacts with the fourth FNIII domain involving aromatic residues Y283 and F1038 **(Figure 2f)**. Notably, these interacting residues are well-conserved across different insect species, suggesting the importance of the capping interactions and conservation of the folded-over conformation (Supplement Figure 4).

The disulfide stapled helical hairpin structure of the N-terminal cysteine rich domain of dROS1 is unusual. However, it reminded us of a structure that we recently solved of the related RTK ALK bound to its ligand ALKAL*^19^*. Indeed, the unique cysteine-rich domain of dROS1 structurally resembles ALK’s ligand ALKAL **(Supplement Figure 5)**. Both share the rare fold of a doubly disulfide-stapled helical hairpin. Therefore, from this prospective, dROS1 could be envisioned as a self-ligated receptor through the N-terminal cysteine rich domain, stabilizing a bent-over conformation.

A Dali*^20, 21^* structural comparison search of the entire cysteine-rich domain (including all four helices) shows the highest structural similarity to the complement component protein C3a **(Supplement Figure 6a, 6b)**. Similar to dROS1’s cysteine-rich domain, C3a also has two helical hairpins. However, the disulfide bond connections are largely not maintained **(Supplement Figure 6c, 6d)**. The disulfide bond connecting helix-3 (H3) and helix-4 (H4) is the only one that is structurally conserved between C3a and dROS1. Whereas, the other disulfide bonds in C3a are inter-hairpin rather than intra-hairpin links. Notably, in the structure of the unprocessed complement C3 (PDB:2A73)*^22^*, the C3a peptide interacts with the complex via the loop connecting H3-H4 **(Supplement Figure 6e)**. This is equivalent to capping loop 2 in the cysteine-rich domain of dROS1, suggesting a possible functional preservation of this loop. In addition, C3a binds to and activates that C3a receptor (C3aR) – a GPCR*^23, 24^*. Interestingly, the ligand of dROS1 (BOSS) is also a GPCR. Therefore, given that dROS1’s N-terminal cysteine-rich domain structurally mimics a GPCR ligand, this finding suggests that crosstalk between dROS1 and BOSS might be bidirectional. As the ECR of BOSS functions as a ligand for dROS1, so too the ECR of dROS1 might function as a ligand for BOSS. Importantly, BOSS is also an orphan GPCR with an as yet to be identified ligand. Given the uniqueness of this cysteine-rich domain and its role in stabilizing the dROS1 N-terminal ECR, we designate it as a CATCH domain for **C**3**a**-**T**ype **C**ysteine-rich **H**elical domain.

### Structural prediction of BOSS reveals a “Venus-flytrap” architecture similar to mGluRs

The evolution of BOSS diverged so rapidly that tracing homologues by sequence is incredibly difficult*^25^*. However, the sequence of BOSS’ transmembrane segments suggest homology to metabotropic glutamate receptors (mGluRs) among Class-C GPCRs*^25^*. An AlphaFold*^26-28^* prediction of BOSS’ ECR suggests it adopts a single domain bi-lobed “Venus-Flytrap” (VFT) architecture also similar to mGluRs **(Figure 3a, 3b).** A Dali *^20, 21^*structural comparison search of the predicted structure shows the highest similarity to mGluR-2 **(Figure 3c; Supplement Figure 7)**. Based on structural alignment with mGluR-2, BOSS’ ECR reveals several unique features. This includes an N-terminal loop, an additional C-terminal strand, and a unique C-terminal loop **(Figure 3c, 3d)**. The disulfide bond holding the N-terminal loop in BOSS is also not conserved in mGluR-2 **(Figure 3a, 3c)**.

**Fig. 3.**
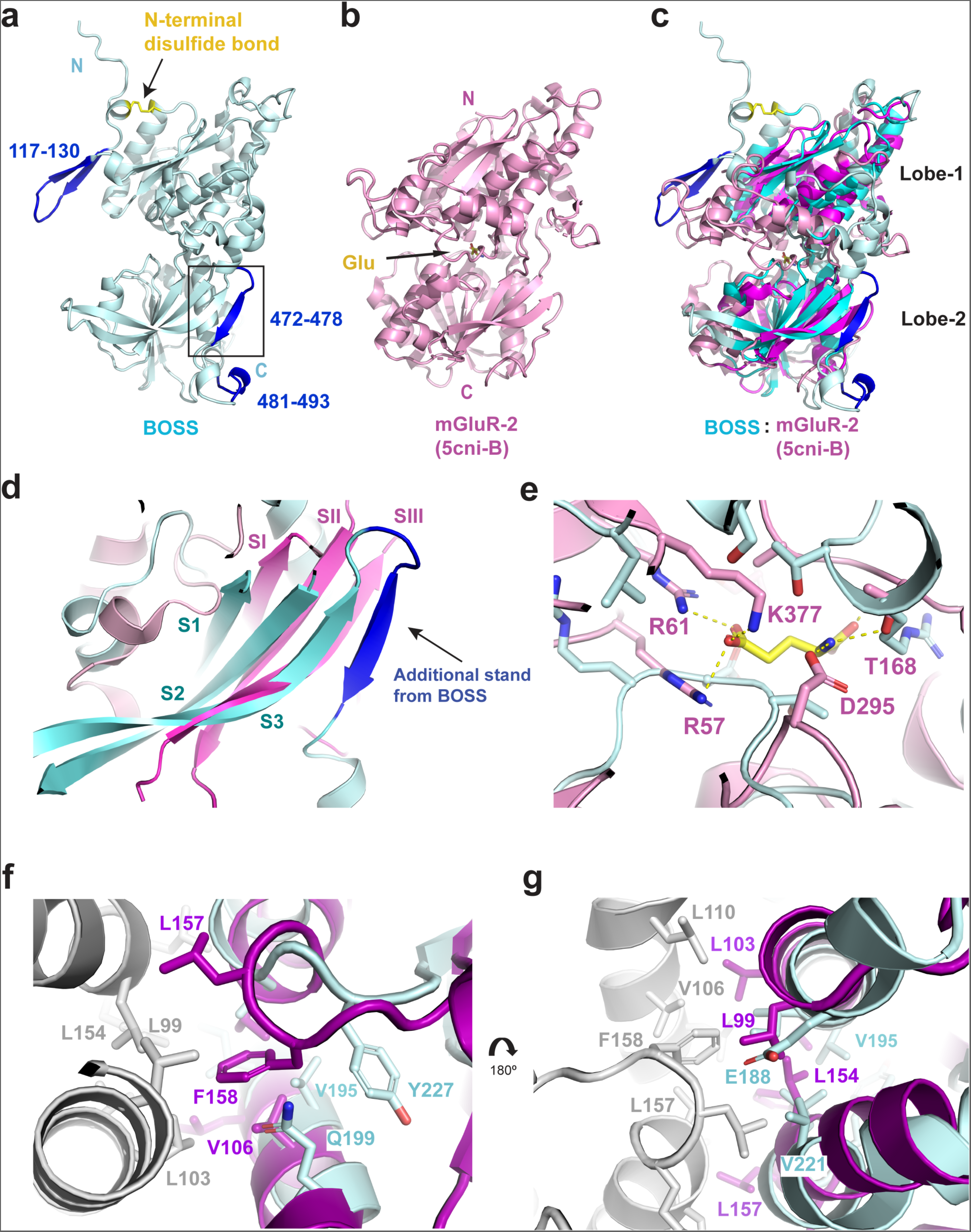
BOSS’ ECD structure is predicted to have a Venus-flytrap fold similar to mGluRs. **a**, Top-ranked model of BOSS ECR generated from AlphaFold colored in light cyan. Unique N-terminal loop and C-terminal loop of BOSS compared with mGluR-2 are shown in blue. **b**, Crystal structure of mGluR-2 (PDB ID: 5cni)*^44^* colored in pink. **c**, Structural alignment of BOSS’ ECR with mGluR-2 generated from Dali server (RMSD=5.2). Aligned region is highlighted in cartoon. **d**, Detailed look at the C terminal unique strand from BOSS (S1, S2, S3: Strand 1, 2 and 3 in BOSS; SI, SII, SIII: Strand 1, 2 and 3 in mGluR). **e**, Key glutamate binding residues from mGluR are not conserved in BOSS. **f**, Alignment of BOSS’ ECR with the full-length inactive dimer of mGluR2 (PDB ID: 7epa; chain A in grey, chain B in purple)*^45^*. The key hydrophobic residue F158*^46^* is not conserved in BOSS’ ECR. Other hydrophobic residues mediating the homodimer in mGluR2 (V106, L157) are either hydrophilic (Q199 in BOSS) or not oriented in the dimer interface. **g**, Other view of the alignment in **f**. Residue L99 in mGluR2 is hydrophilic in BOSS (E188). The hydrophobic residues in BOSS (V221, V295) are not oriented in the dimer interface.

In order to determine if BOSS’ unique N-terminal loop (residue 117-130) was critical for dROS1 binding we performed mutagenesis that removed the loop. Binding analysis of the N-terminal truncated BOSS (BOSS_Δ117-130_) showed no significant changes in binding compared to the wild type BOSS **(Supplement Figure 8a, 8b)**. This indicates that the unique N-terminal loop is not responsible for dROS1 binding. Notably, while BOSS shows structural similarity to mGluRs, it does not seem to retain the conserved residues involved in glutamate binding found in mammalian mGluRs **(Figure 3e)**. Additionally, the residues that dictate dimerization of mGluRs are also not conserved in BOSS **(Figure 3f, 3g)**. This indicates that BOSS’ function is distinct from that of mGluRs. To further examine the potential effect of glutamate on the BOSS-dROS1 interaction, we performed binding studies in the presence and absence of 2 mM L-glutamate. The results show that adding glutamate does not affect the binding affinity **(Supplement Figure 8c)**. Thus, the interaction between dROS1 and BOSS is not regulated by the presence of glutamate.

### Mapping of binding interfaces in the dROS1-BOSS complex using HDX-MS

To find the binding interface most critical for the interaction between dROS1 and BOSS, we used hydrogen-deuterium exchange and mass spectrometry (HDX-MS). HDX-MS probes structural flexibility by the exchange of the amide hydrogen atoms of the protein backbone with deuterium*^29^*. By comparing deuterium uptake of peptide regions between each monomeric protein and the complex (ι1uptake), potential binding regions can be identified. We have previously used this technique to map the binding epitope of antibodies targeting the RTK ALK*^19^*. For this experiment we used the dROS1 (1-4) construct and the full-length BOSS ECR. These constructs were stable at the concentrations necessary for the experiment and also include the interaction sites **(Figure 1c)**. Given that the affinity between these proteins is relatively weak (∼ 3-5 µM K_D_), we developed a strategy to preserve the interaction by immobilizing his-tagged protein on Ni-NTA resin to increase the local concentration (details in **Material and Method** section).

The deuterium uptake of different regions is compared and ranked to identify regions that show statistically significant protection from exchange after complex formation (**Figure 4**, also see **Material and Method** section for details). To aid in visualization, regions protected from exchange (i.e. stabilized regions) were mapped onto the structures of dROS1 (**Figure 2b**) and BOSS (**Figure 3a**). HDX-MS results show that one of the top protected regions in BOSS upon dROS1 binding includes the C-terminal beta-strand “VTNVTTY” (residue 472-478, **Figure 4a, 4b, blue**). Other stabilized regions in BOSS include a part of an alpha-helix (residue 365-370) in the core of BOSS’ structure, a short alpha-helix (residue 221-226), and a short peptide in the N-terminal loop (residue 114-118) **(Figure 4a, 4b, grey)**. In addition, the binding of dROS1 to BOSS enhances structural disorder in a central beta-strand in the second lobe (residue 445-463) in the vicinity of the C-terminal beta-strand “VTNVTTY” **(Supplement Figure 9)**.

**Fig. 4.**
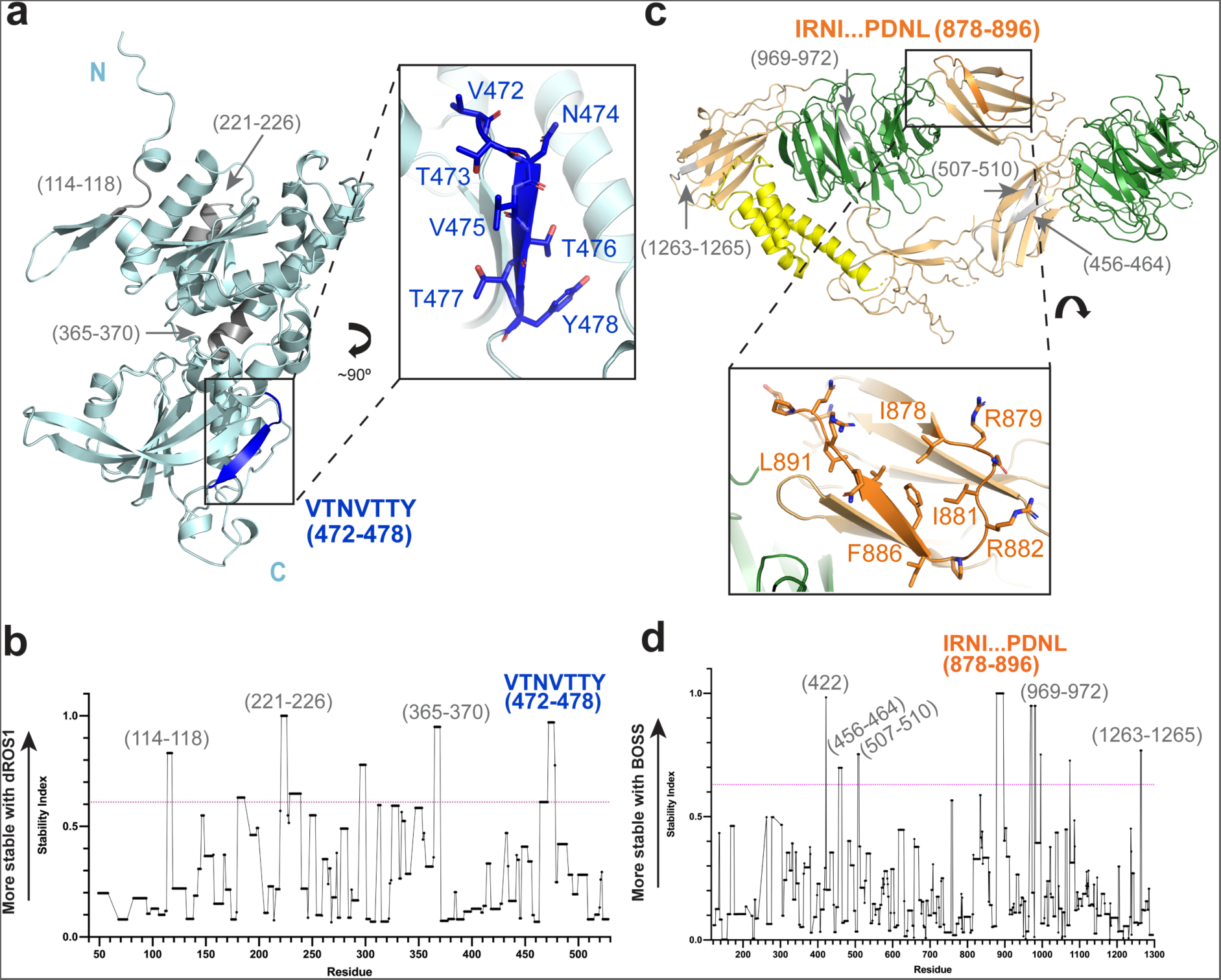
HDX analysis of dROS1 binding to BOSS. **a**, Potential dROS1 binding region in BOSS (VTNTTY) is mapped on an AlphaFold predicted BOSS structure (in blue) with a closer view shown (inset). **b**, His-tagged BOSS binding to dROS1 was probed by HDX-MS as described in the Materials and Methods. The stability index was plotted against the residue number of BOSS. The magenta dotted line indicates statistical significance (ι1uptake >0.61 Da at 85 % confidence based on Student’s t-test). Regions with the stability index above the magenta dotted line show statistically significant stabilization upon dROS1 binding, and their residue numbers are indicated. Two independent HDX-MS experiments (n=2 biological repeats) were performed with 3 technical repeats for each n experiment. **c**, Potential BOSS binding region in dROS1 (IR…DNL) is mapped on dROS1’s structure (in orange) with a closer view shown (inset). **d**, Stabilized region of dROS1 upon BOSS binding. The stability index was plotted against the residue number of dROS1. The magenta dotted line shows the stability index at 90 % confidence (corresponds to ι1uptake >0.63Da) based on Student’s t-test. Regions with the stability index above the confidence line show statistically significant stabilization, and their residue numbers are indicated. Two independent HDX-MS experiments (n=2 biological repeats) were performed with 3 technical repeats for each n experiment.

When mapped on the structure of BOSS (**Figure 4a**), peptide 365-370 is not surface exposed and unlikely to directly participate in dROS1. Whereas, peptide 114-118 is part of the N-terminal loop that we previously confirmed does not contribute to dROS1 binding **(Supplement Figure 8b)**. Thus, based on our HDX results aided with structural prediction, peptide 472-478 (C-terminal beta-strand “VTNVTTY”) may represent a direct binding epitope for dROS1.

Similarly, stabilized regions in dROS1 upon BOSS binding are found in peptides spanning residues 456-464 and 507-510 in the second FNIII domain **(Figure 4c, 4d, grey)**, and 878-896 in third FNIII domain **(Figure 4c, 4d, orange)**. These regions are within the construct which we determined to be important for binding **(Figure 1c)**. Within these regions, peptide 878-896 in the third FNIII domain is more surface exposed and thus is more likely to be a direct interaction site. We also found another stabilized peptide (residue 969-972) located outside of the binding region, which likely represents an indirect stabilization of protein dynamics upon binding **(Figure 4c, 4d, grey)**. Together, our HDX data combined with our structure and binding data narrows down the likely complex binding epitope to a strand in dROS1’s third FNIII domain and a strand near BOSS’ ECR C-terminal loop.

### Structural prediction of the dROS1-BOSS complex

In the absence of a dROS1-BOSS complex structure, we generated a theoretical model of the complex using AlphaFold*^26-28^*(see **Material and Methods**). Since our binding studies narrowed down the binding epitope to the N-terminal region of dROS1, we were able to apply this constraint when generating models. Multiple predicted complex structures all converge to the same binding epitope between dROS1 and BOSS. Importantly, the predicted structures are all consistent with our HDX-MS and binding experiments **(Figure 5a)**.

**Fig. 5.**
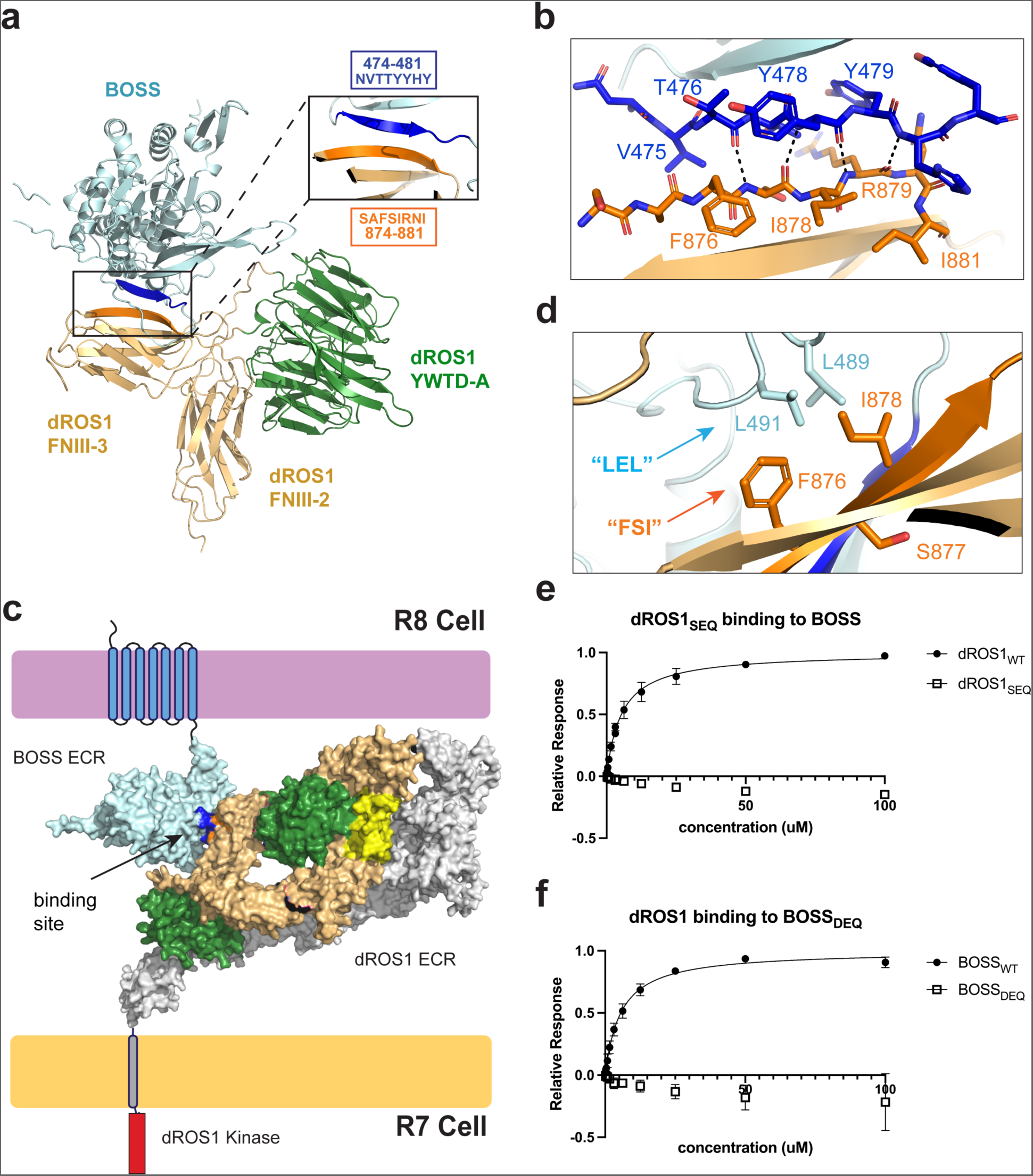
Structural model of the complex between dROS1 and BOSS. **a**, AlphaFold prediction of dROS1 (construct 2-3) in complex with BOSS’ ECR. The predicted model shows the interaction site is located at the third FNIII domain (FNIII-3) of dROS1 and the C-terminal loop of BOSS’ ECR, and mediated though parallel beta-strand augmentation. **b**, A detailed view of the parallel beta-strand augmentation is shown including F876 and I878. **c**, Model of dROS1 and BOSS binding at the cell surface. Cyan: BOSS’ ECR; yellow: CATCH domain; light orange: N-terminal dROS1 FNIII domains (1-4); green: dROS1’s YWTD-A and YWTD-B; grey: model of the remaining ECR domains generated from AlphaFold. Binding site is indicated with blue for BOSS and orange for dROS1. **d**, BOSS’ hydrophobic Leucine residues (L489 and L491) interact with the hydrophobic core of dROS1’s third FNIII domain. **e**, Binding analysis shows dROS1_SEQ_ (mutation of F876 and I878) does not interact with BOSS (n=2 for both dROS1_WT_ and dROS1_SEQ_). **f**, Binding result shows that the “LEL” to ‘DEQ” mutation in BOSS disrupts binding to dROS1 (construct 1-9) (n=3 for BOSS_DEQ_, n=2 for BOSS_WT_).

In these models we found that the complex interaction is mediated by either “parallel” or “anti-parallel” beta-strand augmentation between dROS1’s third FNIII domain (874-881) and a C-terminal strand of BOSS’ ECR (474-481) **(Figure 5a, 5b; Supplement Figure 10a, 10b)** – the very epitopes identified in our HDX-MS. By placing the two complex models between opposing cell membranes, we find that the parallel model orients better than the anti-parallel model – which generates a clash between the receptor and the membrane **(Figure 5c; Supplement Figure 10d, 10e, 11)**. Thus, we favor the parallel model.

We wished to test the predicted complex model using mutagenesis. However, directed mutagenesis – attempting to disrupt the complex interaction – is difficult to generate as the interaction is mediated by main chain strand augmentation rather than side chains. Whereas, mutation that overtly disrupts the strand would likely lead to misfolding of the domain itself. Although, in addition to the main chain beta-strand augmentation, a hydrophobic interaction is predicted in the parallel model to be mediated by surface exposed Leucine residues (L489 and L491) in BOSS **(Figure 5d)**. These Leucine side chains are modeled to interact with the hydrophobic core of the third FNIII domain in dROS1 (FSI, 876-878) **(Figure 5d)**. Notably, the “FSI” residues are not conserved between the second and third FNIII domains in dROS1, which could explain the specificity for the interaction with the third FNIII domain **(Supplement Figure 8d)**. Rather than the hydrophobic residues “FSI”, the second FNIII domain has the sequence “SEQ”. Therefore, in an attempt to disrupt the complex interaction by mutagenesis without disrupting the fold, we mutated the “FSI” of third FNIII domain into the corresponding residues “SEQ” found in the non-binding second FNIII domain. Binding results of dROS1_SEQ_ demonstrate no observable binding to BOSS **(Figure 5e)**. This result is consistent with the model, however, the dROS1_SEQ_ construct was less stable and less pure than the WT construct.

To further validate the model, we sought to mutate the surface exposed hydrophobic residues on BOSS that are predicted to interact with the FSI of dROS1’s third FNIII domain – the reciprocal to the dROS1_SEQ_ experiment. Notably, the Leucine residues (L489 and L491) are located at BOSS’ ECR C-terminal loop that is structurally distinct and not found in mGluRs) **(Figure 3a)**. Mutation of Leucine residues in BOSS to non-hydrophobic residues “LEL” to “DEQ” (BOSS_DEQ_) expressed as well as the WT indicating that the mutations did not affect protein stability. SPR analysis of BOSS_DEQ_ binding to dROS1 (Construct 1-9) shows no detectable binding. This finding is additionally consistent with the parallel model of the interaction between dROS1 and BOSS **(Figure 5d, 5f)**.

## Discussion

dROS1 is homologous to the human RTK and oncogene ROS1, which is the least well-studied RTK*^15^*. Multiple ROS1 fusion proteins have been discovered to play important role in a variety of human cancers including glioblastoma, non-small-cell lung cancer, and breast cancer*^30, 31^*. Due in part to the difficulty of purifying ROS1, no structural insight has yet been established to reveal the architecture of its ECR region. In addition, ROS1 is the last orphan human RTK, with its ligand remaining to be firmly established. Also, a mammalian BOSS equivalent has yet to be identified*_10_*. Thus, ROS1 is likely similar to the RTK ALK, where the ligand controlling its activation is not structurally conserved between invertebrates and vertebrates*^32-35^*.

We find here that dROS1 is regulated by relatively weak interactions with BOSS that are enhanced by the high local concentration when restricted to two dimensions between juxtaposed cell membranes. Our biophysical characterization narrows down the binding epitope between the interaction of dROS1 and BOSS. This could provide insight into the ligand binding region of mammalian ROS1. Indeed, for ALK, despite having structurally different ligands for different species, the binding regions are largely maintained*^19^*. Alternatively, the structural similarity of BOSS with mGluRs indicates that a possible regulatory ligand of mammalian ROS1 might be an mGluR or another class C GPCR.

Earlier efforts on elucidating the dROS1 activation mechanism showed that neither soluble BOSS’ ECR nor dimerized/oligomerized BOSS’ ECR is able to stimulate dROS1’s kinase activity*^9, 36^*. Instead, all 7 transmembrane helices of BOSS are required to induce R7 differentiation*^36^*. In our studies, we show that the N-terminal CATCH domain is structurally similar to the protein C3a. Intriguingly, the C3a receptor (C3aR) has a small ECR and C3a binds deeply into the TM region of the receptor*^24^*. This points to a very intriguing possibility, that the CATCH domain might interact with the TM region of BOSS and this interaction may help regulate the activation of dROS1. Importantly, our studies were unable to directly test this hypothesis because our binding studies were done with the isolated ECR of BOSS and did not include the TM domains.

Here we show the first glimpse of the structure of a ROS1 receptor and gain insight into how it is regulated by ligand. An early model of dROS1 synthesis and processing proposed that cleavage of the poly-arginine sequence yields an ECR non-covalently associated with the kinase domain*^12^*. Removing the ECR might be essential for receptor activation as the truncated dROS1 is constitutively active*^17^*. Our structure of dROS1’s N-terminal ECR – being compact rather than linear, suggests that the bulky ECR could physically keep the kinase domain at distance and prevent oligomerization and signaling in the absence of ligand binding.

Future studies will focus on investigations with dROS1’s full length receptor to determine whether dROS1 resembles other well-known RTKs and cytokine receptors in their self-assembly upon ligand binding. In addition, more research should be done to determine whether the crosstalk between dROS1 and BOSS is bidirectional, and whether dROS1 regulates BOSS’ functions as a glucose sensor*^37^*.

## Materials and Methods

### Recombinant protein expression and purification

BOSS ECR (102-510) and dROS1 with various constructs as dROS1 1-9 (126-2113), 1-8 (126-1992), 1-6 (126-1795), 1-4 (126-1298), 1-3 (126-931), 2-3 (440-931), were incorporated into pFastBac plasmid (Invitrogen) with octa-histidine tag combined with a Factor Xa cut site at the N-terminus and Flag-tag at the C-terminus. The protein was then expressed in Trichoplusia ni (Hi5) cells driven by baculovirus infection at a density of 2×106 cells/ml. Mutants of BOSS and dROS1 were generated by site-directed mutagenesis using QuickChange Kit (Agilent Technologies). Proteins were purified from the medium (Ex-Cell405, Sigma-Aldrich) 2 days post-infection by directly flowing over Ni-Penta™ Agarose-Base Resin (Marvelgent Biosciences). Resins were washed 3 times with 35 mL of 20 mM imidazole (pH 8.0), 100 mM NaCl, and proteins were eluted first with 18 mL of 200 mM imidazole (pH 8.0), 100 mM NaCl and then with 18 mL of 500 mM imidazole (pH 8.0), 100 mM NaCl. Eluted proteins were further purified by size-exclusion chromatography (HiLoad 26/600 superdex 200 pg, GE Healthcare Life Sciences) in 20 mM HEPES (pH 7.5), 150 mM NaCl. Corresponding peaks were checked by SDS-Page gel and concentrated by Amicon Ultra-4 10kDa MWCO concentrator (Millipore).

The His-tags of BOSS proteins and the mutant were cleaved by Factor Xa protease (New England Biolab) with 1 mM CaCl2 and flowed over Ni-Penta™ Agarose-Base Resin (Marvelgent Biosciences). The flow-through was further purified by size-exclusion chromatography (Superdex 200 Increase 10/300 GL size exclusion column GE Healthcare Life Sciences) in 20 mM HEPES (pH 7.5), 150 mM NaCl. dROS1 (1-4) without his-tag was generated following the same protocol (for the HDX-MS experiment).

Purified protein dROS1 (1-8) was coeluted with insect cell ferritin. To remove the ferritin, the protein was further purified by binding to ANTI-FLAG M2 Magnetic Beads (Sigma Aldrich) and eluted with 3× FLAG Peptide (Sigma Aldrich). The eluted protein was then dialyzed using D-Tube Dialyzers (MWCO 6–8 kDa) to remove the flag peptide.

### CryoEM sample preparation and data collection

For cryo-electron microscopy, 3.5 µl of purified dROS1 (1-8) at 500nM were applied on glow-discharged Quantifoil Cu R1.2/1.3 300 mesh grids, and blotted for 5 seconds at 100% humidity and 16 °C. Samples were plunge-frozen into liquid ethane using Vitrobot Mark IV (FEI) and transferred into liquid nitrogen. Grids were imaged under the 300 KeV Titan Krios electron microscope (FEI) with a K3 summit direct electron detector (Gatan). Images were recorded under super-resolution mode at a pixel size of 0.825 Å/pixel using SerialEM v3.8.6 and Digital Micrograph v3.31.2359.03. Data was collected with a dose rate of 17.5 e-/pix/second for 0.06 s/frame. A total of 7835 images were recorded and processed using CryoSPARC. Detailed parameters of data collection are summarized in **Supplement Table 1**.

### CryoEM data processing and 3D reconstruction

Raw movies were motion-corrected using patch motion correction and contrast transfer function (CTF) parameters were estimated using patch CTF. Bad micrographs were discarded by manual inspection. A small set of particles were manually picked, extracted and 2D classified as templates for autopicking. The template-picked particles were extracted with a box size of 786 Å and Fourier cropped to 192 Å, which was then used to generate 2D classes. Particles with good 2D classifications were re-cropped to 384 Å and re-extracted, which were then used for one round of ab initio reconstruction into 4 classes and iterative rounds of heterogeneous refinement into 5 classes. The best model (388,364 particles) was selected for non-uniform refinement. The overall resolution is 3.74Å estimated by Fourier shell correlation (FSC) at 0.143 cutoff. These particle sets (388,364 particles) were then used in additional rounds of heterogeneous refinement into 4 classes, which gave us a 3.94 Å map (144,311 particles) with more density. These volumes were assessed by local resolution estimate using the CryoSPARC program and the final resolution was colored using UCSF ChimeraX. Both volumes were used in model building. Local mask was created covering the CATCH domain, FNIII-1, FNIII-2 and FNIII-4, from the 3.94 Å map using UCSF ChimeraX and imported back into CryoSPARC. Local refinement was performed using created local mask and yield a slightly more density around the masked area.

### CryoEM model building

The initial model of the structure was generated using AlphaFold2*^26-28^*. The model was segmented into individual domains and fitted into a map using ChimeraX. Given that the 3.74 Å map has a higher resolution, FNIII-1 to YWTD-B were fitted using this map. The cysteine-rich domain and FNIII-4 were fitted into the 3.94 Å map. Domains were then merged and imported into Coot 0.9.6*^38^* from SBGrid*^39^*along with the two maps for manual inspection. Realspace refinement was done using Phenix 1.21-5207*^40^* from SBGrid*^39^*. Detailed statistics and parameters for model building and refinement are provided in **Supplement Table 1**.

### Surface plasmon resonance

Surface plasmon resonance (SPR) experiments of ligand binding were all performed using BIAcore T200 in 20mM HEPES (pH7.5), 100 mM NaCl. The CM5 biosensor chip was activated with N-hydroxysuccinimide (NHS) and N-ethyl-N’-[3-(diethylamino)propyl] carbodiimide (EDC). dROS1 constructs at 30 ng/µL in 10 mM acetate pH 4.0-6.0 were flowed over the activated surface at 10 µL/min for 300 sec and then the activation was ended by 1 M ethanolamine pH 8.5. Purified ligand BOSS proteins at varied concentrations were flowed over the dROS1 surfaces at 10µL/min for 180 sec. The response unit (RU) values corresponding to the plateau, corrected by background binding through subtraction of signal obtained in control surface, were taken as the measure of binding. The binding affinities were calculated by plotting the plateau RU against BOSS concentrations and fit into single-site specific binding model using Prism9. All binding curves were then plotted as BOSS concentrations against maximal binding (Bmax).

### Bio-Layer Interferometry

Experiments were performed using the Octet RED96e instrument. His-tagged dROS1 protein (construct 2-3) were loaded onto penta-His sensors (Sartorius) to a response unit of about 0.5 nm followed by quenching in 10 ng/mL biocytin (Sigma-Aldrich) for 60 sec. The sensors were then equilibrated in HBS buffer (20 mM HEPES, 150 mM NaCl pH 7.5) for 30 sec and dipped in 4.8uM of BOSS_WT_ or BOSS_Δ117-130_, or BOSSWT with 2 mM L-glutamate, for 60 sec, then dissociated in HBS buffer for 60 sec. Background binding of BOSS to the sensor was measured by repeating the same experiment without loading dROS1. The response curves were aligned to the starting point of association, corrected by background binding, and the values corresponding to the plateau of the mutant were plotted relative wild type using Prism9 (n=3 for BOSS_WT_, n=3 for BOSS_Δ117-130_, n=4 for BOSS with 2 mM L-glutamate).

### AlphaFold prediction

dROS1 model was searched from AlphaFold protein structure database (DeepMind and EMBL-EBI) and the construct 1-6 (126-1795) was used to generate the model for CryoEM map. BOSS ECR (102-510) was used for generating single protein prediction. BOSS ECR (102-510) and dROS1 construct 2-3 (440-931) were used for heterocomplex prediction. Modeling of BOSS ECR and BOSS-dROS1 complex was performed using the publicly available ColabFold v1.5.3*^41^*. MSA options were set to ‘mmseqs2_uniref_env’ and pair mode was ‘unpaired_paired’. Anti-parallel complex model was generated without template while the parallel complex was generated with dROS1 2-3 construct from dROS1 model as template. For BOSS ECR, five models were generated and ranked by pLDDT scores (Rank_1 = 84.2, Rank_2 = 83.4, Rank_3 = 83.4, Rank_4 = 82.6, Rank_5 = 81.7). Rank_1 model was used in this study. For parallel complex, five models were generated and ranked by ‘multimer’ matric. Rank_1 model with BOSS C-terminal loop interacting with dROS1 FNIII-3 was selected in this study. For anti-parallel complex, five models were generated and ranked by ‘multimer’ matric. Rank_1 model with the highest ipTM score (Rank_1 = 0.7, Rank_2 = 0.319, Rank_3 = 0.207, Rank_4 = 0.14, Rank_5 = 0.134) was selected in this study. Detailed statistics and figures are provided in **Supplement Figure 12**. Human ROS1 model was searched from AlphaFold protein structure database (DeepMind and EMBL-EBI).

### Hydrogen-Deuterium Exchange Mass spectrometry (HDX-MS) sample preparation

Probing low affinity interaction such as dROS1-BOSS poses a major challenge in HDX-MS experiments because dROS1-BOSS complex dilution with the deuterium labeling buffer (20 mM HEPES, 100 mM NaCl, pD 7.4) leads to the complex dissociation. To maintain at least 80 % of the complex after 10-fold dilution with the deuterium buffer, dROS1 at 0.6 mg/mL (4.5 μM) must be pre-incubated with a large excess amount of BOSS (210 μM, 11.8 mg/mL). Although such an experiment enables one to probe the complex, resulting mass spectra would be largely dominated by BOSS peptides due to the greater amount of BOSS, making dROS1 deuterium uptake assessment extremely challenging. Furthermore, protein sample overloading onto pepsin and UPLC columns can result in column clogging. To circumvent these problems, we designed HDX-MS experiment based on a previous study where the effect of weak EGFR kinase dimerization on the kinase activity was studied by immobilizing the his-tagged kinase domain on DOGS Ni-NTA lipid vesicles*^42^*. The immobilization of the his-tagged protein increases its local concentration such that the kinase domain with weak propensity to dimerize could be dimerized (the kinase remains monomeric up to 50 μM).

To minimize dROS1 or BOSS concentration used for the deuterium labeling and to increase local protein concentration, his-tagged dROS1 (construct 1-4) or BOSS (0.6 mg/mL, 100 μL) was immobilized on 50µL packed Ni-Penta™ Agarose-Base Resin (Marvelgent Biosciences) for 10 min. Then unbound protein was flowed through the resin by gentle centrifugation at 30g for 30 sec. Concentrations were checked before and after centrifugation to ensure most of the proteins were still bound to the resin. Either dROS1 or BOSS (0.6 mg/mL, 100 μL) without his-tag was then co-incubated with the Ni-resin to facilitate the complex formation followed by a brief centrifugation (30g for 30 sec) to remove the unbound protein. Concentrations were checked to ensure most of protein was bound to form the complex. Subsequently, the deuterium labeling buffer (50 μL, 20 mM HEPES, 100 mM NaCl, pD 7.4) was added to the Ni-resin and incubated for 10 min before the addition of cold 50μL quench buffer (200 mM glycine, 100 mM TCEP, pH 2.4) to stop the labeling and elute the complex off the Ni-resin. The labeled sample was immediately flash-frozen in liquid N_2_ and stored at -80°C until mass spectrometry analysis. Similarly, control experiments were carried out with either his-tagged dROS1 or BOSS only. For peptide sequencing experiments, H_2_O buffer (20 mM HEPES, 100 mM NaCl, pH 7.4) was used instead of the deuterium buffer.

Frozen samples were thawed quickly and immediately injected onto an Enzymate BEH pepsin column (Waters) maintained at 2°C to digest proteins for 3 min in UPLC solvent A (water + 0.1 % formic acid) at 100 μL/min. Peptic peptides were trapped in BEH C18 pre-column (2.1 x 5 mm, 1.7 μm, Waters), were separated on BEH C18 analytical column (1.0 x 100 mm, 1.7 μm, Waters), and eluted from the analytical column by 5% to 45 % linear solvent B (acetonitrile + 0.1 % formic acid) gradient over 7 min. HD-MS^e^ data were acquired at 300-1200 m/z range using Synapt G2-Si mass spectrometer (Waters) using 0.4 sec scan time in sensitivity ion mobility mode with Leu-Enk as a lock CCS and mass compound. Other instrument parameters were: capillary voltage, 3 kV; cone voltage, 10 V; source offset, 80V; source temperature, 100°C; desolvation temperature, 200°C; cone gas flow, 0 L/Hr; desolvation gas flow, 700 L/Hr; nebulizer gas, 7.0 Bar. The low and high energy collusion ramp was 5 to 10 V and 20 to 50 V, respectively.

### HDX-MS data processing

ProteinLynx Global Server 3.03 (Waters) was used to sequence each peptic peptide, and the deuterium uptake of peptic peptides was evaluated using DynamX 3.0 (Waters), with the minimum intensity cut-off set to 1000, minimum product per amino acid to 0.1, and maximum MH^+^ error set to 10 ppm. The deuterium uptake of both dROS1 and BOSS peptides is the average uptake of two biological samples (n = 2), with 3 technical repeats/n. The average standard deviations (SD) of two biological repeat experiments are: 0.449 Da (his-tagged dROS1-BOSS complex), 0.333 Da (his-tagged dROS1 only), 0.230 Da (his-tagged BOSS-dROS1 complex), and 0.239 Da (his-tagged BOSS only). The average SD of all technical repeats is in the range of 0.0937 Da – 0.131 Da. To identify dynamic changes in dROS1 upon BOSS binding, the deuterium uptake of his-tagged dROS1 in the presence of untagged BOSS was evaluated, and the uptake was compared to that of the his-tagged dROS1 without BOSS for obtaining the difference in the deuterium uptake (ι1uptake). Similarly, to identify structural changes in BOSS upon dROS1 binding, the deuterium uptake of his-tagged BOSS in the presence of untagged dROS1 was assessed, and the uptake was compared to that of his-tagged BOSS without dROS1. The ι1uptake of all peptides was normalized to the highest ι1uptake value, which is defined as the stability index = 1. The stability index of different regions is mapped as described previously*^43^*. Briefly, the stability index of a region is assigned using the index of the shortest peptide in a given region. For overlapping peptides, the stability index of a peptide that retains the overlapping region at its C-terminus is used. DynamX 3.0 (Waters) uses the same strategy to assign the percent exchange of different regions at single amino acid residue level.

## Acknowledgments

We thank Claudio Alarcon, Yansheng Liu, Mark Lemmon, Kate Ferguson and their laboratories for valuable discussions. We additionally thank Yuhong Zuo and Long Han for their expertise in guiding the CryoEM processing.

## Author contributions

D.E.K. designed the overall project. J.Z. and D.E.K. wrote the manuscript with input from all authors. J.Z. and D.E.K. analyzed the structure. J.Z. generated all materials and performed all the experiments assisted by H.L., T.L., and Y.W. Y.T. designed, conducted, and analyzed the HDX experiment.

## Competing interests

The authors declare that they have no competing interests.

**Correspondence and requests for materials** should be addressed to D.E.K.

## Figures

**Supplement Fig. 1.**
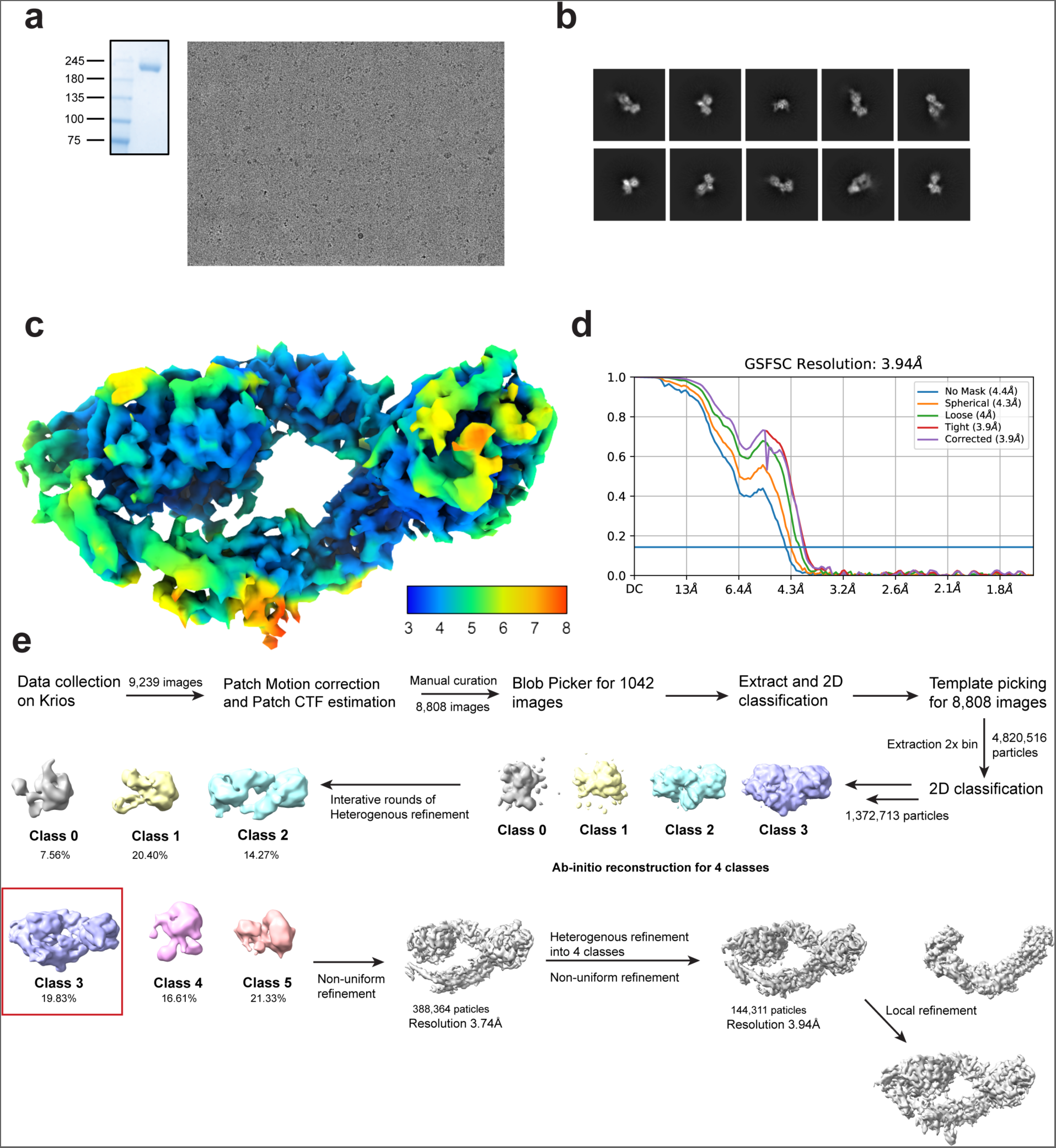
CryoEM map quality, resolution estimate and workflow of data analysis. **a**, SDS-PAGE analysis of protein purification and a representative CryoEM micrograph of dROS1. **b,** Selected 2D class average. **c,** 3D reconstructed map colored with local estimated resolution in angstrom (Å). **d,** Gold-standard FSC curve. **e,** Workflow of data analysis using CryoSPARC4.3.

**Supplement Fig. 2.**
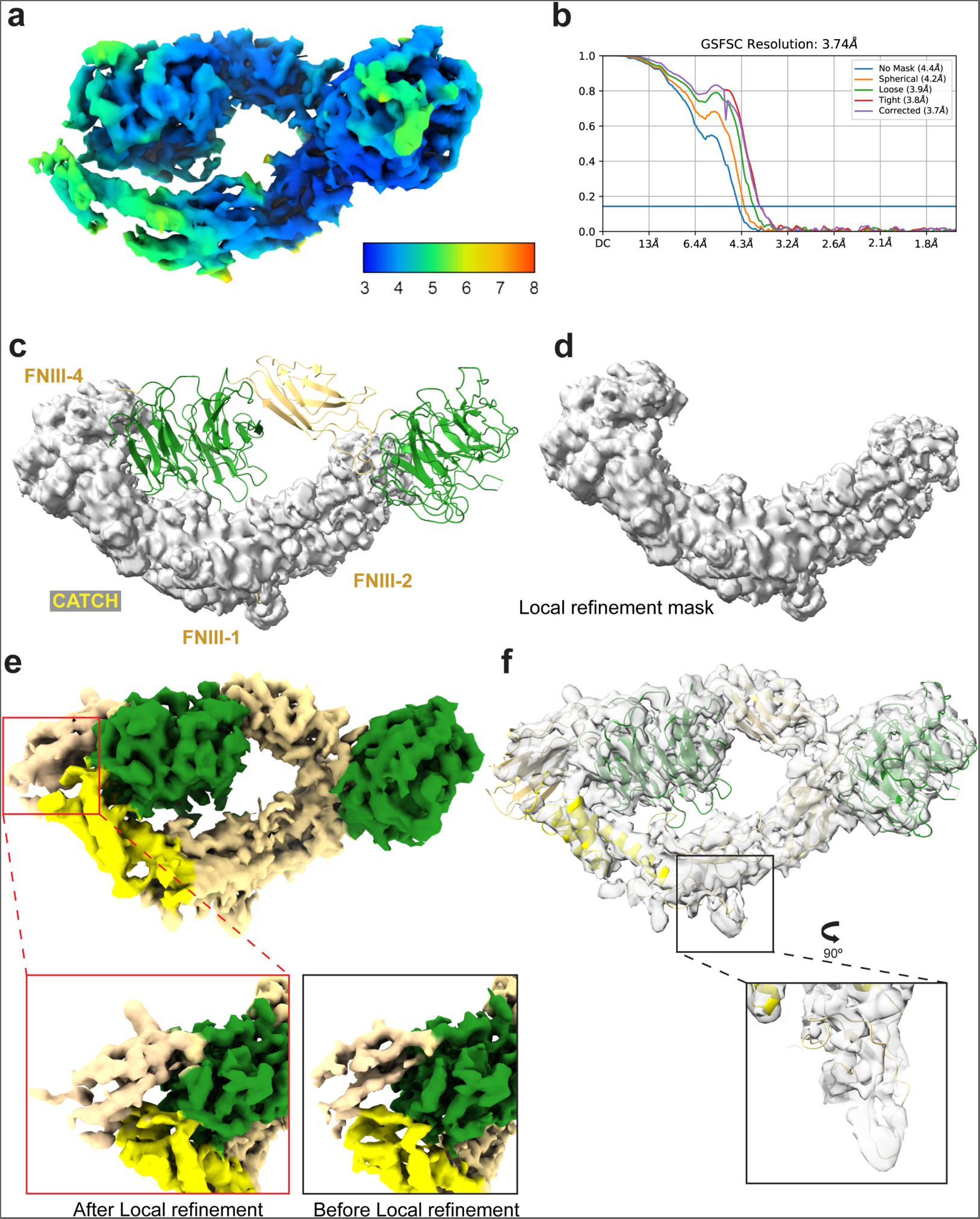
CryoEM map with 3.74 Å and local refinement. **a**, 3D reconstructed map of 3.74 Å resolution from 388,364 particles. Map was resolved with higher resolution but less density near the CATCH domain, thus this map was only used to aid model building. **b,** Gold-standard FSC curve of the 3.74 Å map. **c,** and **d,** Mask creation for local refinement covering the CATCH domain and the FNIII-4. Additional coverage of FNIII-1 and FNIII-2 was used to reach enough volume for local refinement. **e,** Density map after local refinement using CryoSPARC. More density was observed in FNIII-4. However, no side-chain density was found in the capping region. The display level for maps before and after refinement is both at 0.0804. **f,** More density was observed in the loop region within FNIII-1.

**Supplement Fig. 3.**
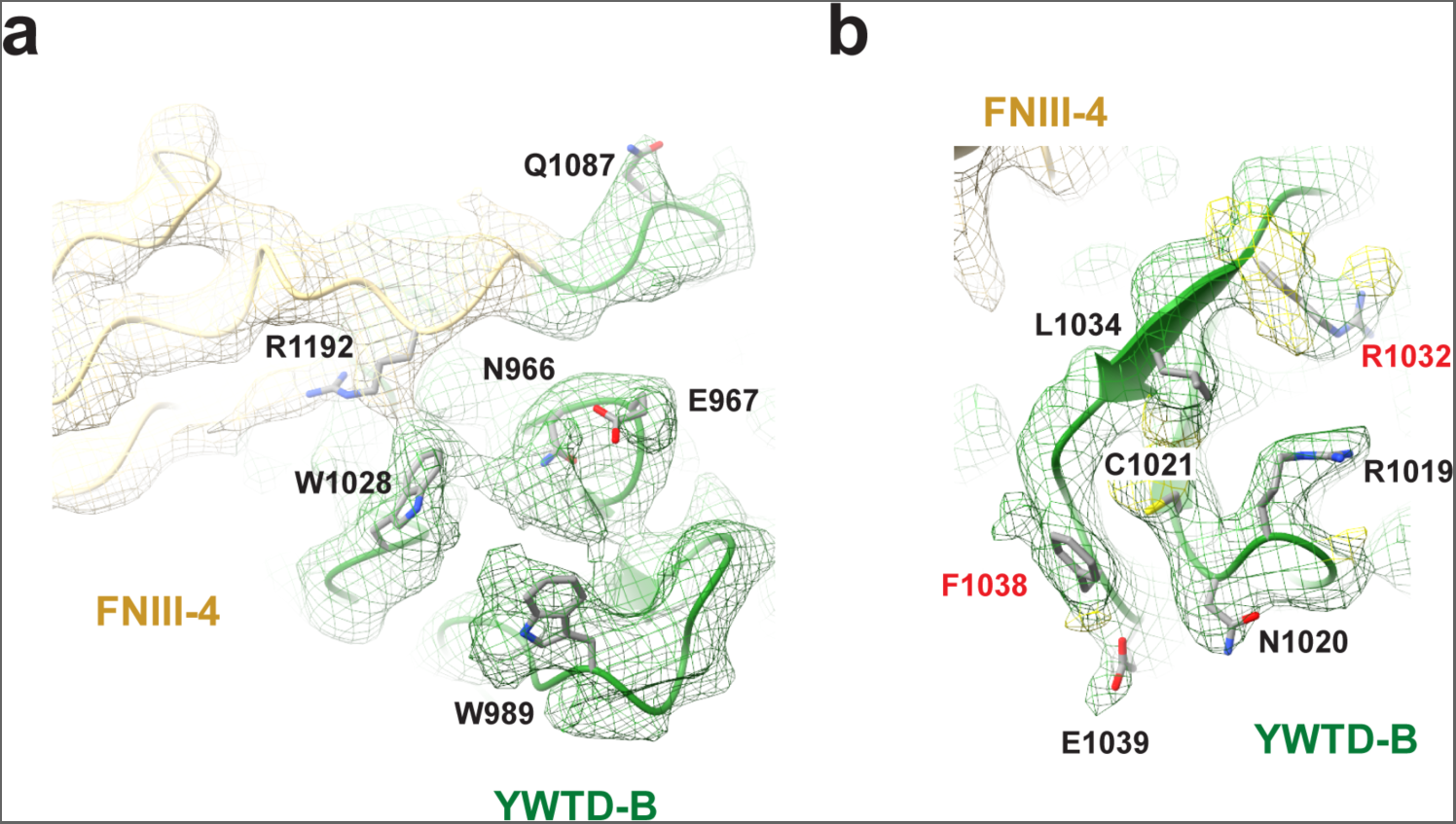
Local density maps of YWTD-B, and FNIII-4. **a** and **b,** EM density maps of YWTD-B and FNIII-4. Residues R1032 and F1038 are participating in the capping interaction (as in Figure 2).

**Supplement Fig. 4.**
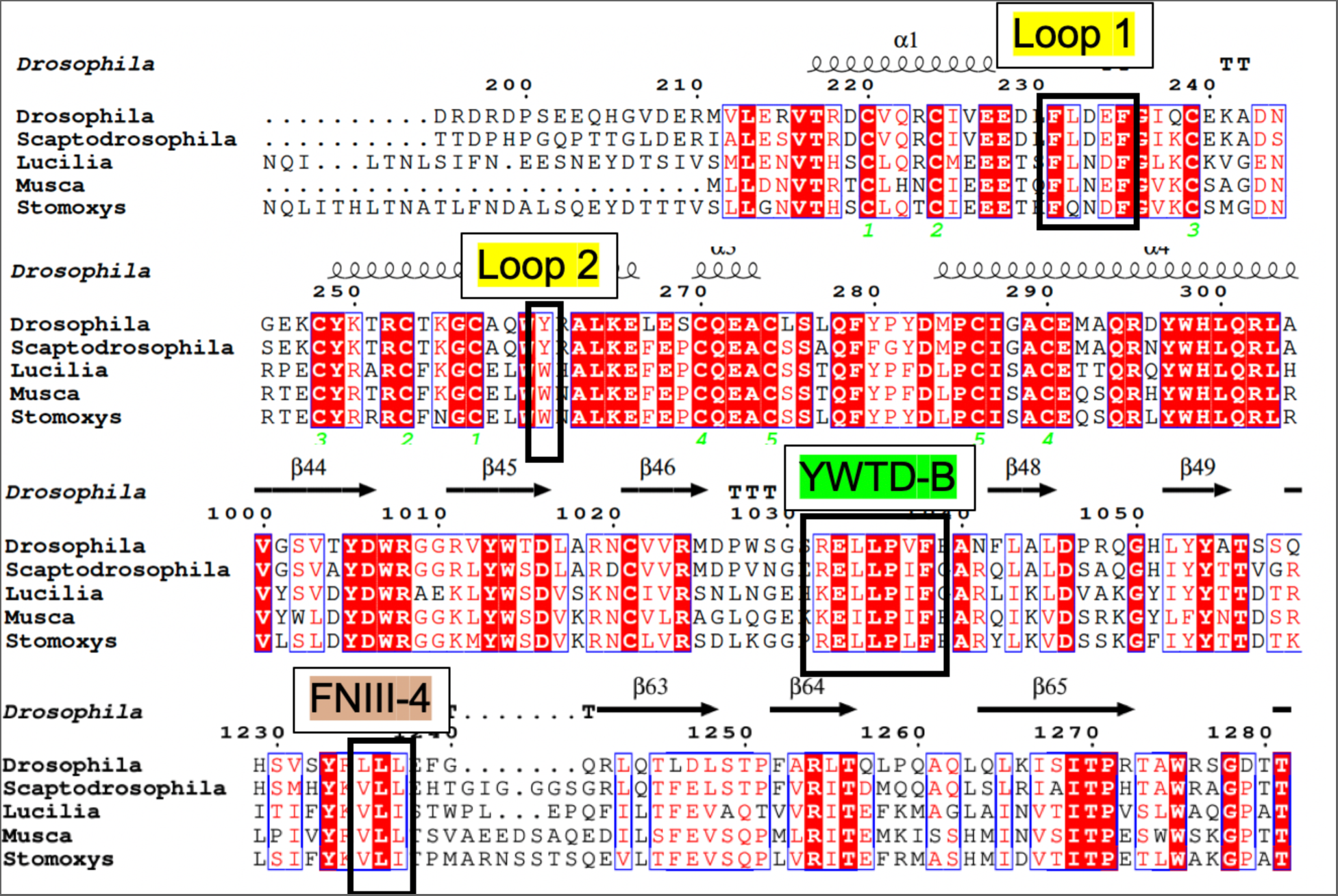
Sequence alignment of CATCH domain through insect species. Residues that are involved in capping interaction were conserved across different insect species. Alignment was performed using pairwise sequence alignment (EMBL-EBI) and ESpript3.0.

**Supplement Fig. 5.**
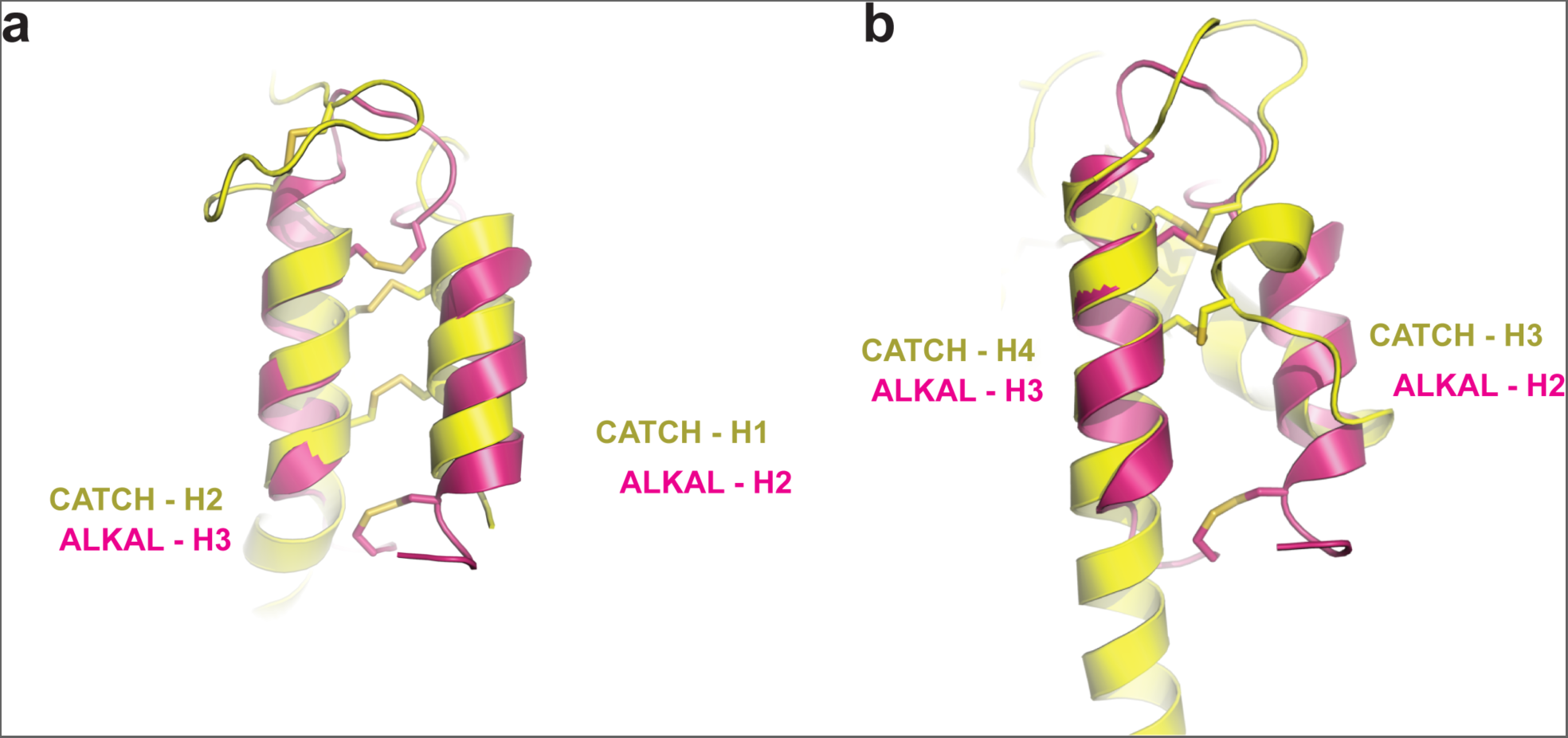
Structural alignment of CATCH domain with ALKAL. **a**, Superimposition of helix-1 and helix-2 (H1, H2 respectively) of the CATCH domain (N-terminal cysteine-rich domain of dROS1) with ALKAL. **b,** Superimposition of the CATCH domain’s H3 and H4 with ALKAL.

**Supplement Fig. 6.**
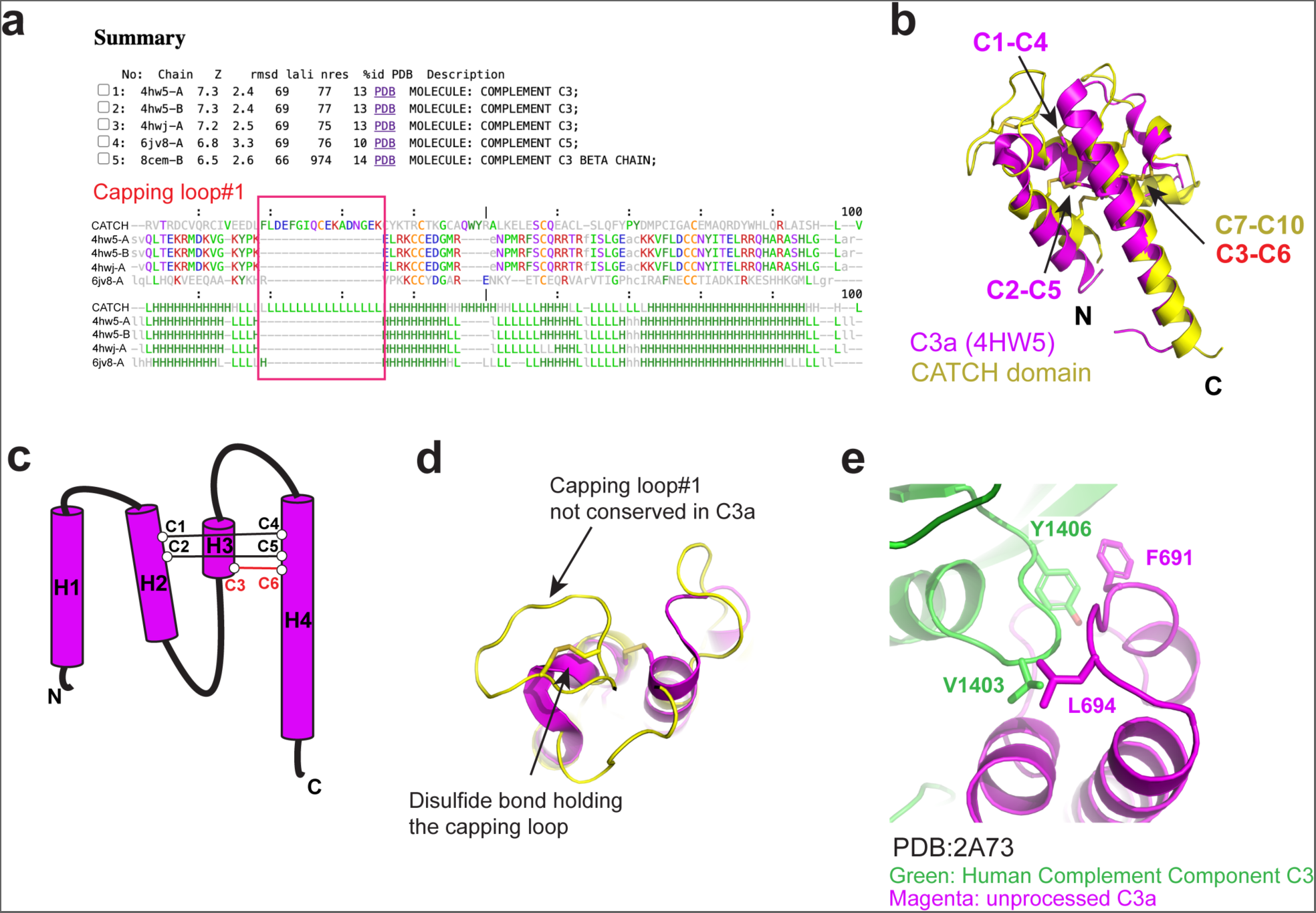
Structural alignment search of the CATCH domain. **a**, Result from 3D alignment search using the Dali server*^20, 21^*. The top candidate was Complement component protein C3a. Secondary structure alignment of the CATCH domain with the top 4 candidates shown. **b,** 3D alignment of CATCH domain with C3a (PDB: 4hw5, chain A)*^47^* (RMSD = 2.4). **c,** Simple illustration of C3a. The protein C3a has 3 disulfide bonds, with C1-C4 and C2-C5 connecting helix-2 (H2) and helix-4 (H4), and C3-C6 connecting H3 and H4. Note that C3-C6 colored in red is located at the same position as C7-C10 in dROS1’s CATCH domain. **d,** Capping loop 1 and the disulfide bond holding the capping loop are unique features of dROS1’s CATCH domain. **e,** In the unprocessed C3 complement protein structure (PDB: 2A73)*^22^*, the loop connecting C3a’s H3 and H4 (equivalent to capping loop 2 in dROS1’s CATCH domain) are interacting with the rest of the protein, similar to the interaction of capping loop 2 of CATCH with dROS1 FNIII-4.

**Supplement Fig. 7.**
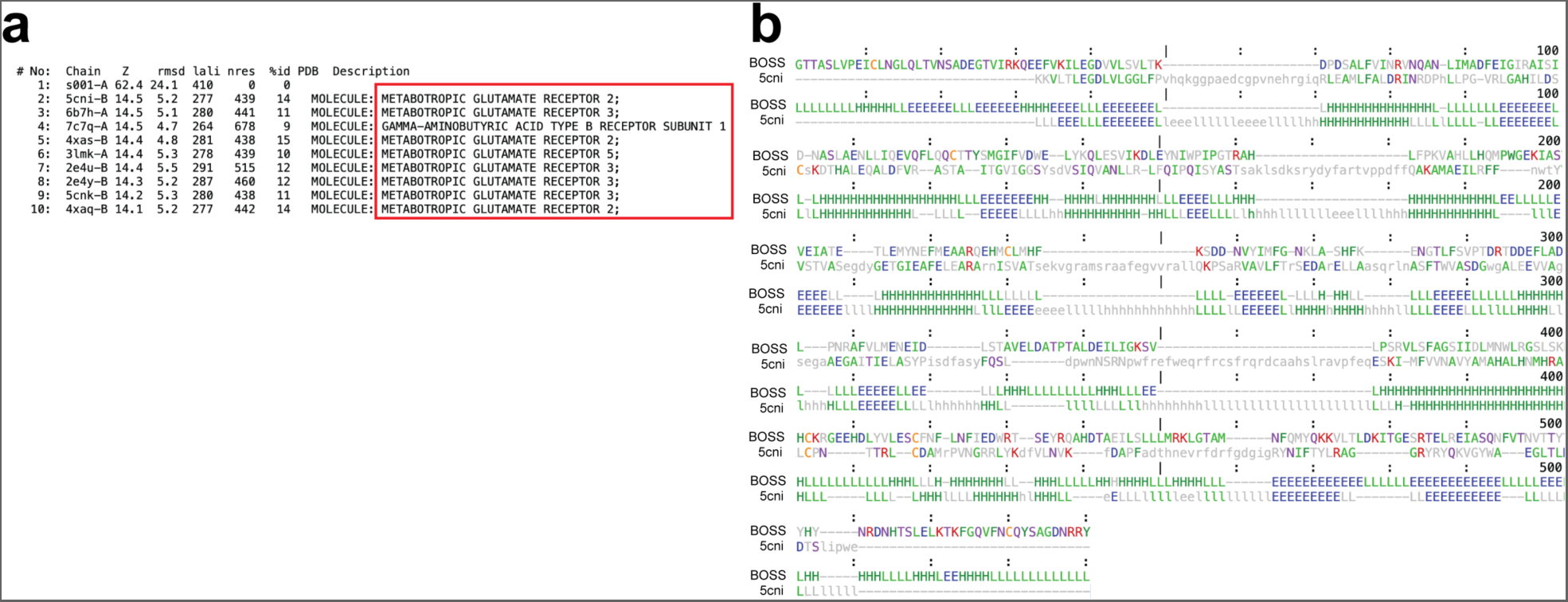
Structural and sequence alignments of BOSS with mGluR. **a**, Summary of 3D structural comparison with BOSS’ ECR as a template using the Dali server*^20, 21^*. Results with the highest Z-scores fall into mGluRs (metabolic glutamate receptors). **b,** Amino acid sequence and secondary structure alignment are shown.

**Supplement Fig. 8.**
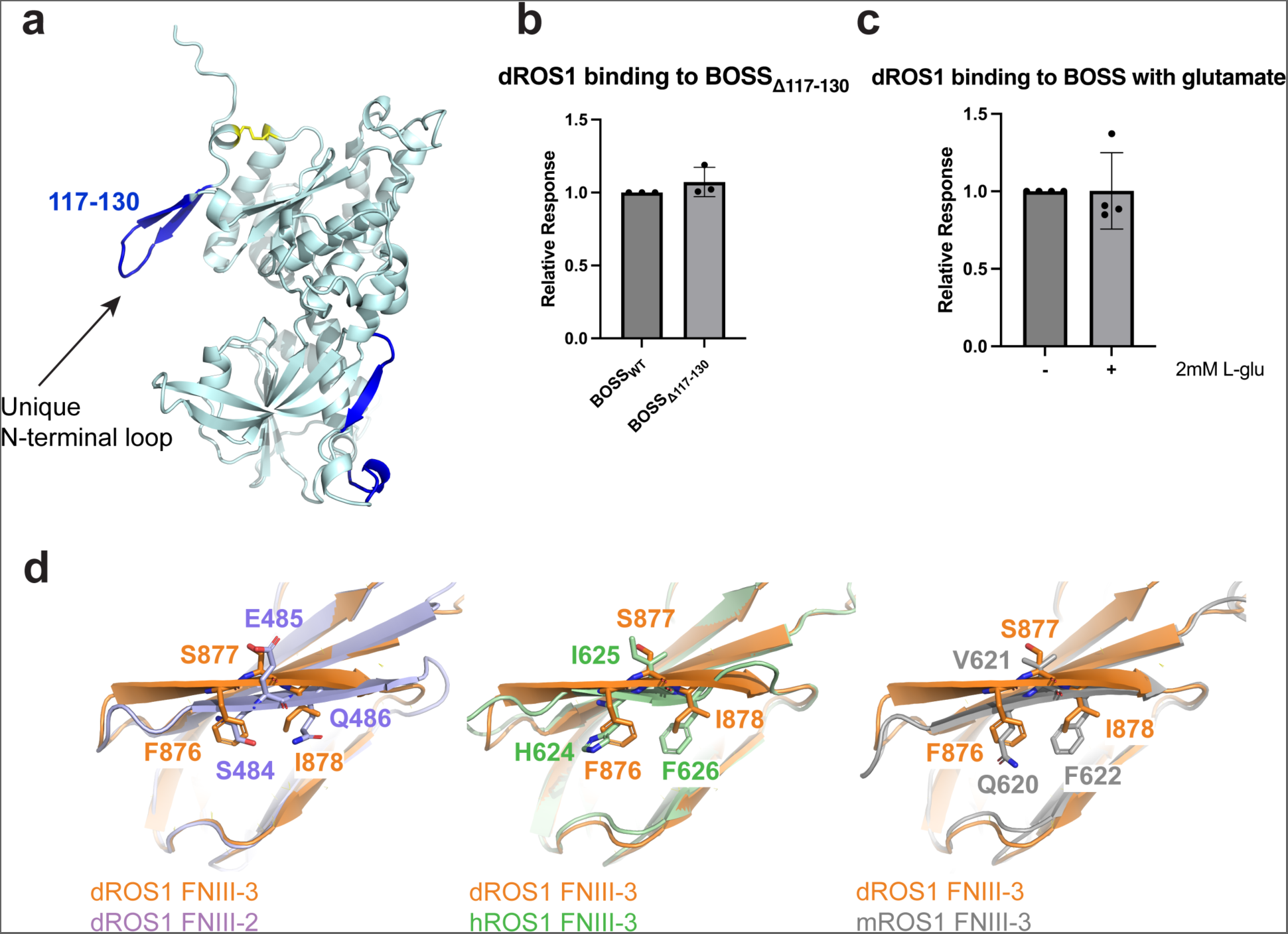
Structural and binding analysis of dROS1 and BOSS mutations. **a**, Unique N-terminal loop of BOSS that is not conserved in mGluRs. **b,** BOSS with truncated N-terminal loop does not affect binding. Unpaired two-tailed Student’s t-tests shows no significant difference between BOSS_WT_ and BOSS_Δ117-130_ (P = 0.2822). **c,** BOSS_WT_-dROS1 interaction is not affected by the presence of glutamate. Unpaired two-tailed Student’s t-tests shows no significant difference (P = 0.9777). **d,** Overlay of dROS1 FNIII-3 with dROS1 FNIII-2, hROS1 (human ROS1) FNIII-3, and mROS1 (mouse ROS1) FNIII-3. the hydrophobic residues “FSI” dROS1 FNIII-3 is not conserved in FNIII-2 (SEQ). In addition, the corresponding residue of F876 is not hydrophobic in hROS1 and mROS1.

**Supplement Fig. 9.**
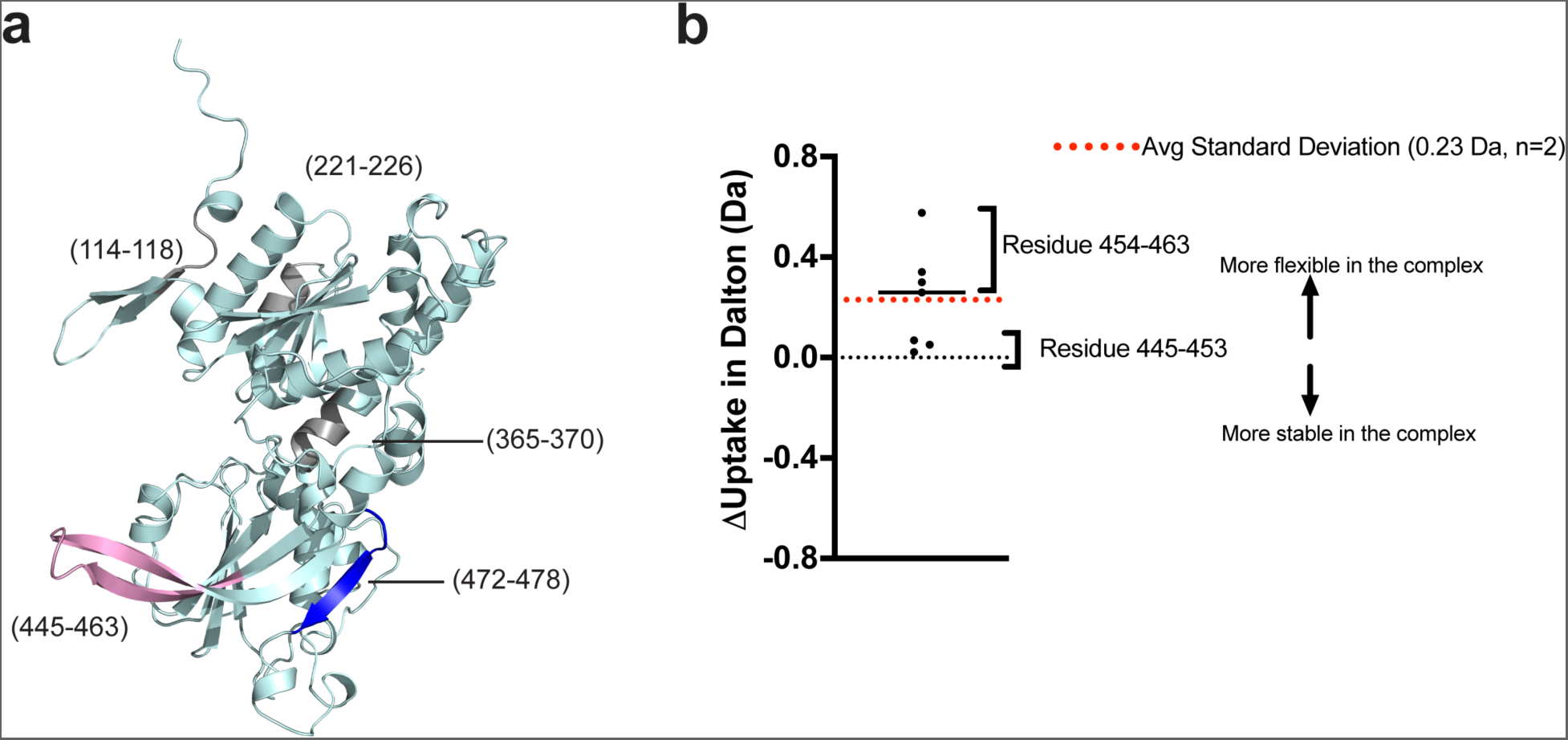
HDX results mapped on the structure of BOSS. **a**, Identified peptides from BOSS mapped in AlphaFold predicted structure. Stable regions were colored in blue or gray with residue numbers indicated. A region colored pink (residue 445-463) demonstrates increased flexibility upon the complex formation. **b,** Δuptake showing residue 445-463 becomes more flexible in complex.

**Supplement Fig. 10.**
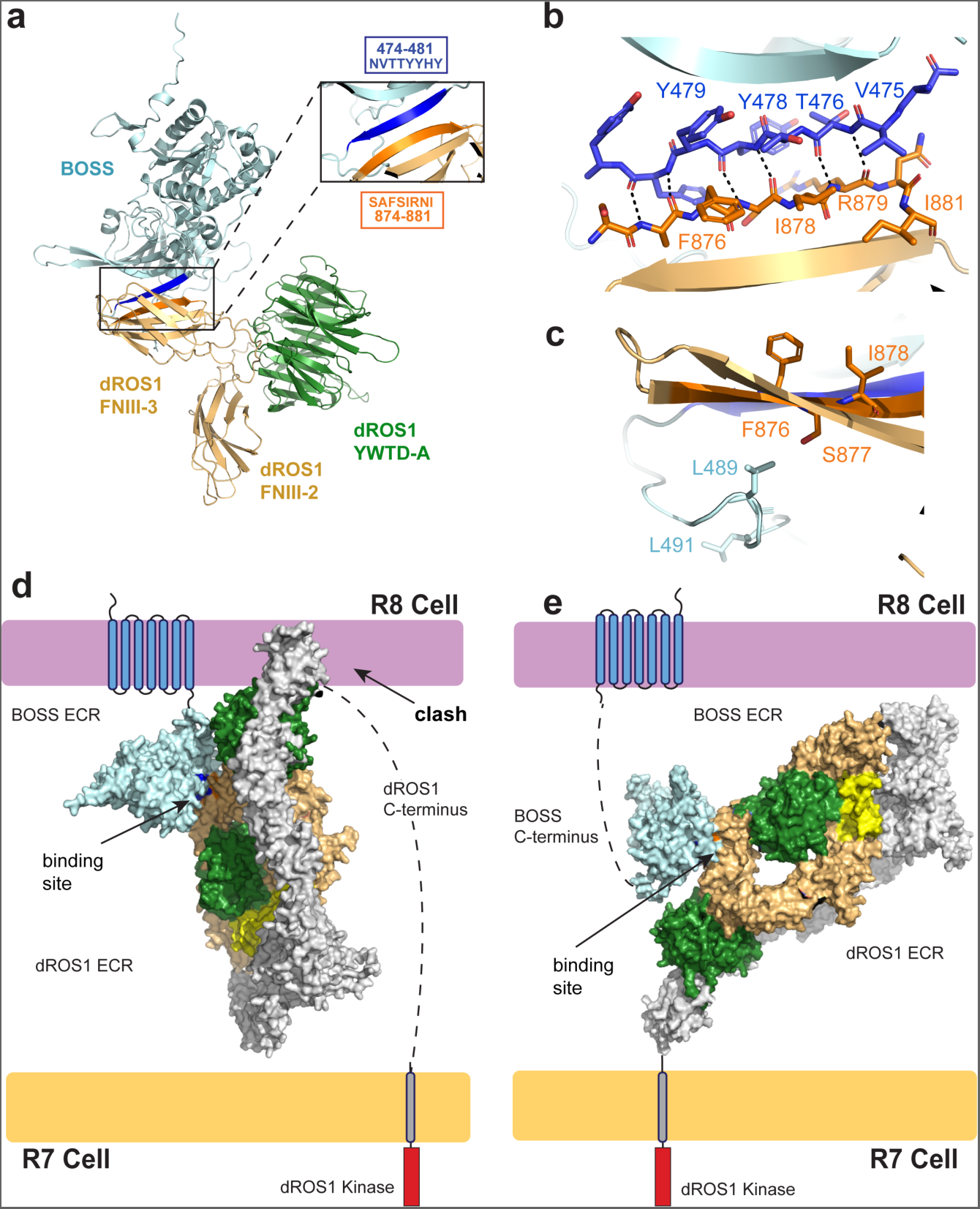
The predicted antiparallel dROS1-BOSS complex structure is incompatible with opposing membranes. **a**, The anti-parallel model of dROS1 in complex with BOSS’ ECR from AlphaFold. While the binding site is consistent with the parallel model shown in Figure 5, the interaction is mediated though “anti-parallel” beta-strand augmentation. **b,** Detailed view of the anti-parallel beta-strand augmentation. **c,** Hydrophobic residues L489 and L491 on BOSS’ ECR C-terminus are positioned outside of dROS1’s third FNIII’s hydrophobic core (not interacting) in the anti-parallel model. **d and e,** The transmembrane regions of BOSS and dROS1 are facing the same direction in the anti-parallel complex model, making it hard to orient the two receptors on opposing cell surfaces. Panel **d** is constrained by BOSS’ TM region with ECR forming a near right angle with the TM as shown in **Supplement** Figure 11. **e,** model constrained by dROS1’s TM region. The dashed lines show the distance that would be required to accommodate the orientation of opposed cell membranes.

**Supplement Fig. 11.**
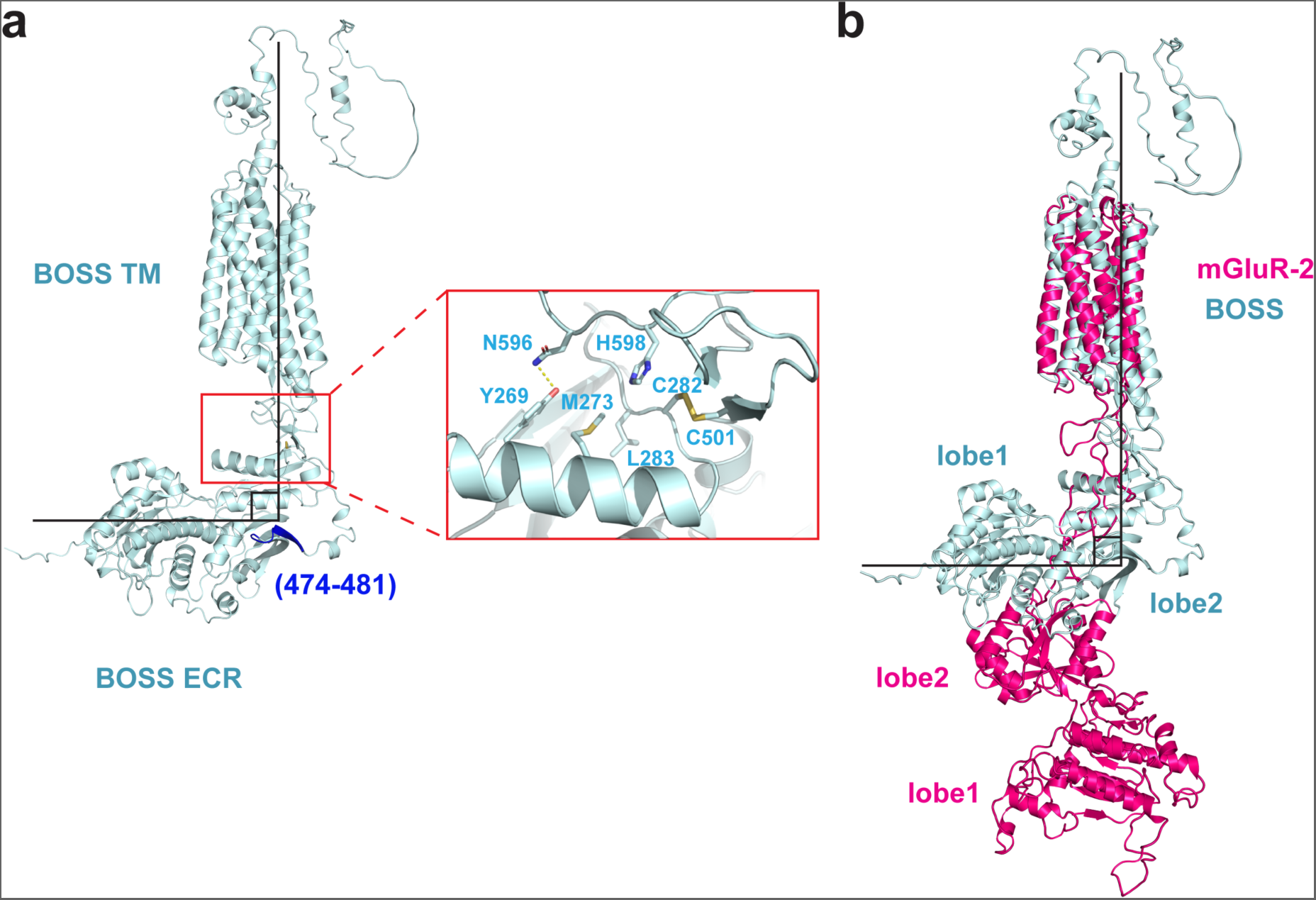
The BOSS ECR is predicted to form a right angle with the TM region. **a**, AlphaFold predicted structure of full-length BOSS. The ECR of BOSS forms a right angle with the TM region, stabilized by hydrogen bonding between N596 and Y269 and constrained by a disulfide bond formed by C282 and C501. This orientation exposes the binding site (highlighted in blue) to the binding of dROS1. **b,** Alignment of the TM regions of BOSS and mGluR-2 (PDB: 7EPA)*^45^* shows this right angle is specific to BOSS.

**Supplement Fig. 12.**
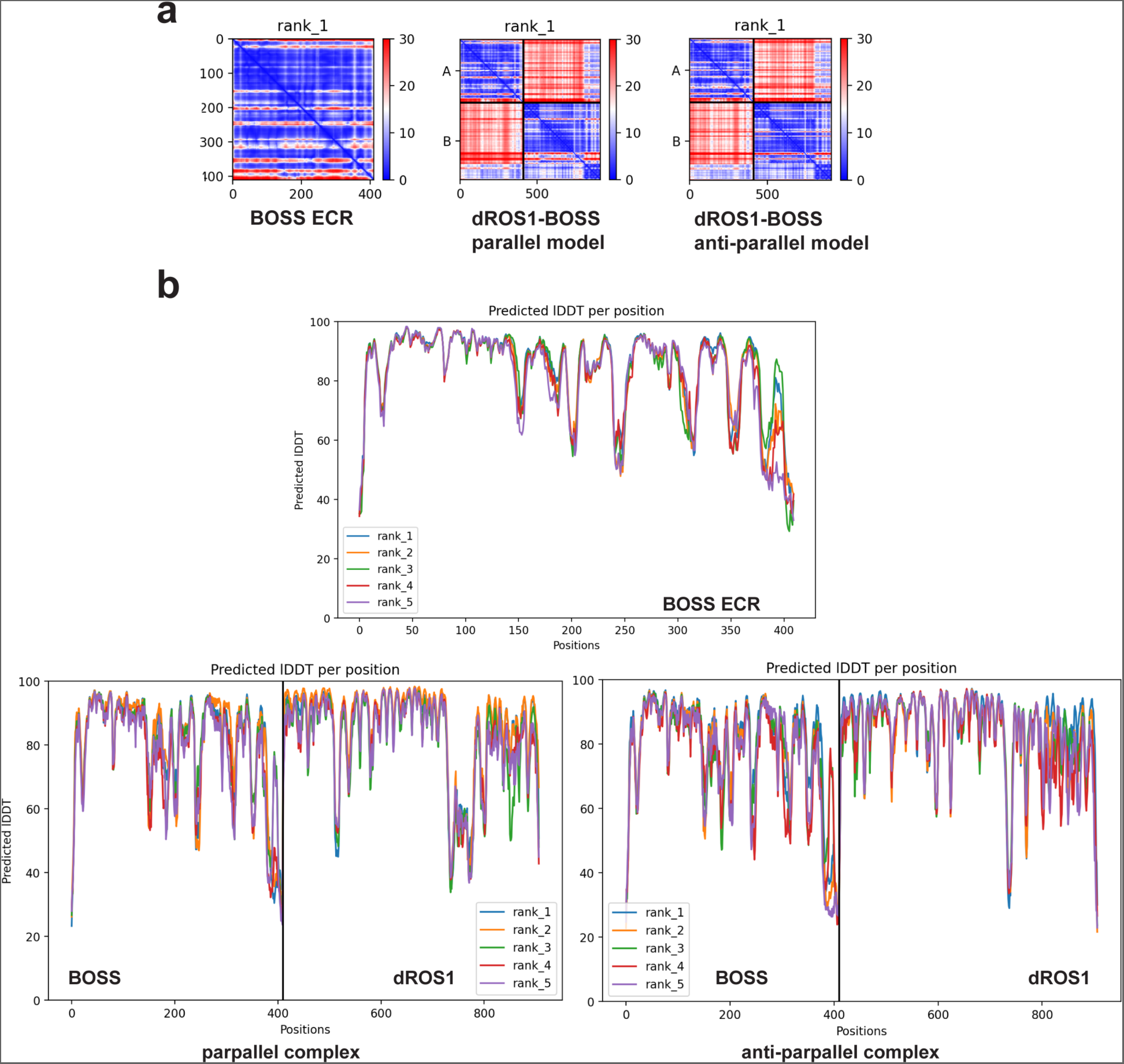
Summary of AlphaFold statistics of BOSS and dROS1-BOSS complex. **a**, Predicted alignment error plots (PAE) for all models. For the heteromeric complex plots, “A” on the Y-axis represents BOSS and “B” represents dROS1. **b,** pLDDT plots for all models.

**Table A.1.** CryoEM data collection and refinement statistics of dROS1 ECR.

|  |  |  |
| --- | --- | --- |
| **Protein** | dROS1 |  |
| PBD ID | / |  |
| **Data Collection** |  |  |
| Microscope | FEI Titan Krios |  |
| Magnification | 105K |  |
| Voltage (keV) | 300 |  |
| Electron exposure (e<sup>-</sup>/ Å<sup>-2</sup>) | 50 |  |
| Detector | Gatan K3 |  |
| Pixel size (Å/pixel) | 0.825 |  |
| Defocus range | -0.8 to -2.0 |  |
| Final number of particles for reconstruction | 144,311 | 388,364 |
| Symmetry imposed | C1 | C1 |
| Box size (pixels) | 768 | 768 |
| Map resolution (Å) | 3.94 | 3.74 |
| **Refinement** |  |  |
| Number of non-H atoms | 8376 |  |
| Protein residues | 1048 |  |
| Molprobity score | 2.24 |  |
| Clashscore | 17.12 |  |
| Rotamer outlier (%) | 0.44 |  |
| r.m.s.d. values |  |  |
| Bond length (Å) | 0.006 |  |
| Bond angles (Å) | 0.865 |  |
| Ramachandran analysis |  |  |
| Favored (%) | 91.46 |  |
| Allowed (%) | 8.25 |  |
| Outliers (%) | 0.29 |  |

